# Belief states and categorical-choice biases determine reward-based learning under perceptual uncertainty

**DOI:** 10.1101/2020.09.18.303495

**Authors:** Rasmus Bruckner, Hauke R. Heekeren, Dirk Ostwald

## Abstract

In natural settings, learning and decision making often takes place under considerable perceptual uncertainty. Here we investigate the computational principles that govern reward-based learning and decision making under perceptual uncertainty about environmental states. Based on an integrated perceptual and economic decision-making task where unobservable states governed the reward contingencies, we analyzed behavioral data of 52 human participants. We formalized perceptual uncertainty with a belief state that expresses the probability of task states based on sensory information. Using several Bayesian and Q-learning agent models, we examined to which degree belief states and categorical-choice biases determine human learning and decision making under perceptual uncertainty. We found that both factors influenced participants’ behavior, which was similarly captured in Bayesian-inference and Q-learning models. Therefore, humans dynamically combine uncertain perceptual and reward information during learning and decision making, but categorical choices substantially modulate this integration. The results suggest that categorical commitments to the most likely state of the environment may generally give rise to categorical biases on learning under uncertainty.

## Introduction

Humans often have to learn reward contingencies under considerable perceptual uncertainty about the current state of the environment (Bach and Dolan, 2012; Ma and Jazayeri, 2014; Rao, 2010). For example, when learning which varieties of wild berries are edible, perceptual uncertainty about the berries can significantly degrade the correct credit assignment between the berries available in the current environment (states, e.g., currant vs. holly berry) and their taste (rewards, e.g., edible vs. non-edible). Especially when the berries’ taste is additionally uncertain due to natural variability, an unexpected outcome can be the consequence of either the reward uncertainty or having eaten the wrong berry due to perceptual uncertainty. While previous work has extensively studied learning in tasks with clear perceptual information (Dreher and Tremblay, 2016; Glimcher and Fehr, 2013; Rangel et al., 2008; Rushworth and Behrens, 2008), it remains elusive how humans learn under perceptual uncertainty (Schutte et al., 2017; Summerfield and Tsetsos, 2012).

The decision-making literature suggests that humans can consider perceptual uncertainty in perceptual-choice tasks to maximize obtained rewards. Several studies that used paradigms with asymmetric rewards for selecting correct options (e.g., a higher reward for choosing correctly on the left versus right side) have shown that humans trade-off uncertainty about the corresponding perceptual evidence and reward magnitudes (Diederich and Busemeyer, 2006; Kivilcim et al., 2018; Leite and Ratcliff, 2011; Mulder et al., 2012; Simen et al., 2009; Summerfield and Koechlin, 2010; Whiteley and Sahani, 2008). However, humans may not always optimally treat perceptual uncertainty, potentially because they categorize perceptual information (Fleming et al., 2013). Participants often commit to one interpretation of a stimulus (e.g., currant), although other interpretations are possible but neglected (holly berry). One potential explanation for these kinds of categorical effects is that perceptual inference is not only based on currently available perceptual information but also previous perceptual choices that exert categorical biases on future decisions (Luu and Stocker, 2018; Stocker and Simoncelli, 2007; Urai et al., 2019).

Normative theories propose that perceptual uncertainty can be formalized with a belief state that expresses the probability of task states conditional on the current sensory information (Daw, 2014; Dayan and Daw, 2008; Russell and Norvig, 2010). When the belief state is clearly in favor of a particular berry variety, the berry can easily be identified based on the available perceptual information. In contrast, when the belief state indicates comparable probabilities of multiple varieties, berries can not be distinguished. If humans consider perceptual uncertainty during learning, they may use the belief state to regulate reward-based learning (e.g., belief-state weighted learning about berry edibility) (Babayan et al., 2018; Lak et al., 2017, 2020; Starkweather et al., 2017). However, if previous categorical perceptual choices bias learning, then learning is not flexibly regulated according to the belief state based on the current sensory information but a “categorical” belief state that depends on the previous perceptual choice (e.g., commitment to one interpretation of the berry variety and categorical learning about edibility).

Bayesian inference offers normative models for how learning should be regulated when observations are uncertain (Barber, 2012; Bishop, 2006; Murphy, 2012). Previously developed models successfully capture human learning in uncertain and changing environments (Behrens et al., 2007; Mathys et al., 2014; Nassar et al., 2010; Payzan-LeNestour et al., 2013; Yu and Dayan, 2005), and it could, therefore, be possible that Bayesian learning based on belief states also captures reward-based learning under perceptual uncertainty. However, an alternative implementation is belief-state Q-learning, where learning based on prediction errors is modulated as a function of the belief state (Chrisman, 1992). Recent work suggests that such algorithms explain key-aspects of learning under perceptual and other forms of state uncertainty in animals (Babayan et al., 2018; Lak et al., 2017, 2020; Starkweather et al., 2017). However, it is currently unclear how learning under perceptual uncertainty differs between normative Bayesian-inference and Q-learning models.

This study aimed to examine how belief states and categorical-choice biases determine human learning under perceptual uncertainty. To formally test this question, we developed several artificial agent models (Hassabis et al., 2017; Russell and Norvig, 2010). In particular, a normative Bayesian belief-state model in which the belief state rendered learning under perceptual uncertainty more cautious to avoid that misinterpreted uncertain perceptual information corrupts learning. We also developed a categorical Bayesian agent that ignored perceptual uncertainty because the prior categorical perceptual choice exclusively determined learning and a mixture model where both belief states and categorical-choice biases were considered. Similarly, we applied a belief-state Q-learning agent that adjusted the learning rate according to the belief state, a categorical Q-learner, and a mixture of the two. The results show that human participants regulate learning and decision making under perceptual uncertainty according to belief states. Crucially, we also identified a categorical-choice bias that led to a less cautious regulation of learning under perceptual uncertainty compared to our normative Bayesian model. The Bayesian mixture model that considered both belief states and categorical-choice biases captured participants’ learning and decision-making behavior better than the Q-learning mixture model. However, primarily because the Q-learning model was more complex, which was penalized in the model comparison. This interplay between belief states and categorical-choice biases may reveal a general tendency of the brain to regulate learning not only according to uncertainty but also categorical commitments to the most likely state of the environment.

## Results

### Experimental paradigm

To examine how humans learn under perceptual uncertainty, we analyzed data of 52 younger adults that completed a novel Gabor patch perceptual choice-augmented bandit task (Figure 1a). The goal in the Gabor-bandit (GB) task was to maximize the amount of collected reward, which required reward-based learning under perceptual uncertainty. The task consisted of three stages. The first stage required a perceptual decision based on the contrast difference between two Gabor patches, which varied across trials and induced perceptual uncertainty. Half of the participants were asked to indicate the high-contrast patch; the other half was asked to indicate the low-contrast patch. The second stage required an economic choice between two fractals based on a comparison of the fractals’ expected values. Finally, in the third stage, the task delivered a probabilistic reward.

**Figure 1.**
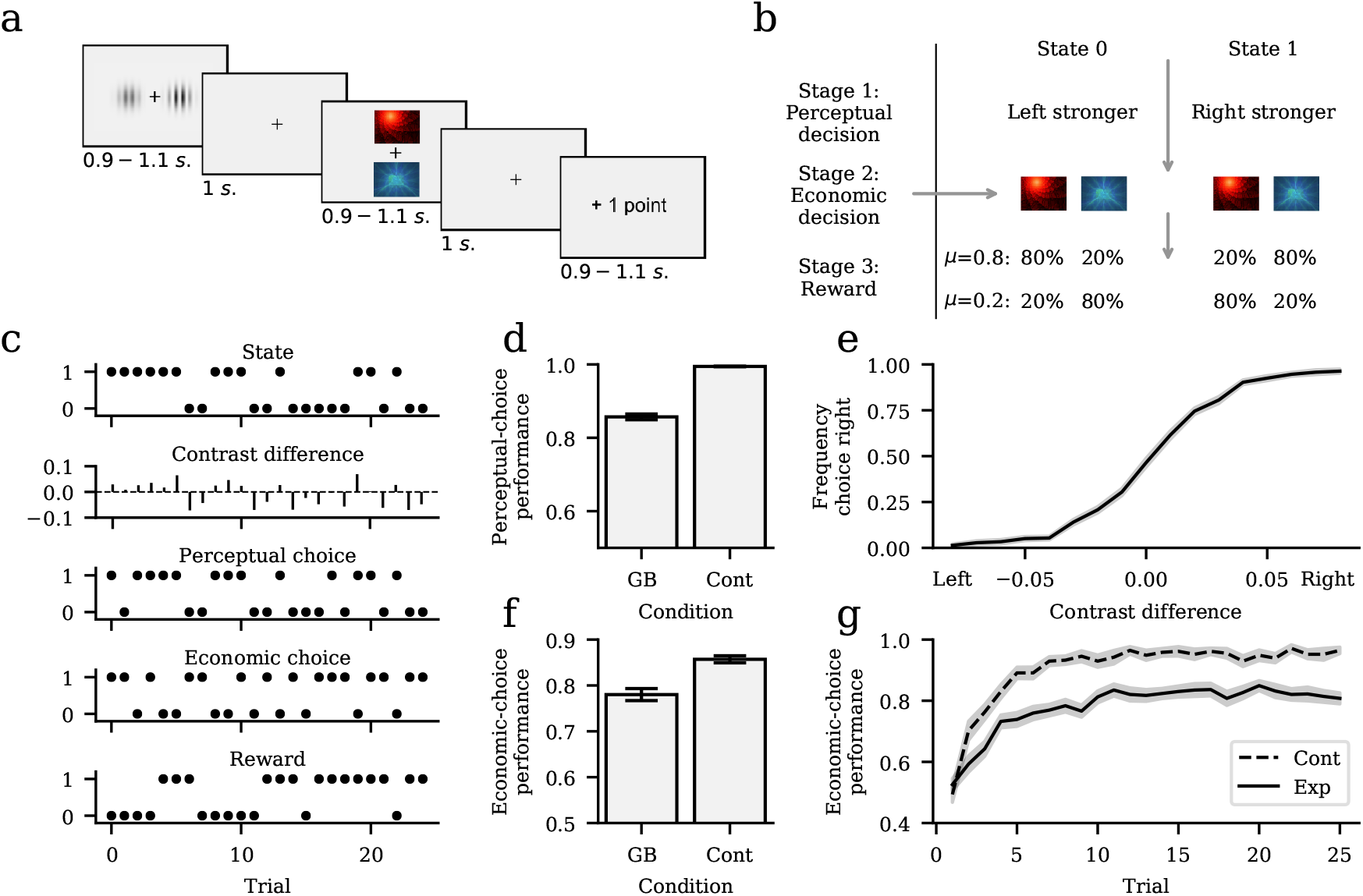
Experimental Task and choice performance. **a)** The Gabor-bandit (GB) task is a state-action-reward contingency learning task that combines perceptual and economic decision-making aspects. Participants were trained to learn state-dependent (Gabor patches) associations between actions (fractals) and rewards (0 or 1 point) over each block of 25 trials, of which a single trial is depicted in the figure. In the first stage, participants reported which Gabor patch displays a higher contrast (perceptual decision). In the second stage, they chose between the red and blue fractal (economic decision). In the third stage, they received a probabilistic reward in response to the economic choice. **b)** Task structure. In each block, the contingency parameter *μ* governs the state-action-reward contingencies. During each trial, the GB task can assume either state 0 or 1. The state determines the position of the high-contrast Gabor patch and the fractal reward probabilities. In state 0, the left patch’s contrast is stronger than the contrast of the right patch. When the contingency parameter *μ* = 0.8, then the red fractal choice option has a higher reward probability (80%) than the blue fractal (20%). In state 1, the right patch contrast is stronger, and the blue fractal has a higher reward probability. In the other half of the blocks, the contingency parameter was *μ* = 0.2, yielding the opposite outcome contingencies. Please also note that the participants’ perceptual decisions had no consequences for the reward delivery. **c)** Illustration of an example participant during a task block. Trial-by-trial task states were randomly 0 or 1 and determined the contrast difference between the left and right Gabor patch (negative difference indicates a stronger contrast on the left side). During the observation of the Gabor patches, the participant made perceptual decisions about the position of the high-contrast patch. Subsequently, during the observation of the fractals, the participant reported the economic decision about the high-reward fractal, which was followed by a probabilistic reward of either 0 or 1 point. **d)** We applied the GB task and a control experiment without perceptual uncertainty where contrast differences were clearly discriminable. Bar plots show mean ± standard error of the mean (SEM) perceptual-choice performance in the GB experiment and the control experiment (Cont). **e)** Perceptual choice psychometric function of the GB experiment indicating a higher frequency of perceptual decisions in favor of the right Gabor patch when the right patch was more clearly the high-contrast patch. **f)** Mean ± SEM trial-by-trial economic-choice performance in the GB and control experiment. **g)** Economic-choice performance (frequency of high-reward fractal choices) in the GB and control experiment increased as a function of trials throughout a block.

As shown in Figure 1b, the task could assume two hidden states (states 0 and 1), which determined the contrast difference between the Gabor patches and the reward probabilities of the fractals. In state 0, the left patch displayed a stronger contrast than the right patch, and in state 1, the right patch had a stronger contrast. Depending on the task state and the contingency parameter that had to be learned throughout a block, the red or blue fractal had a higher reward probability. In half of the blocks, the contingency parameter *μ* was 0.8, indicating that in state 0, the red fractal had an 80% reward probability and the blue fractal a 20% reward probability. In state 1, the reward contingency was reversed, that is, the red fractal had a 20% reward probability and the blue fractal 80%. In the other half of the blocks, the contingency parameter was equal to 0.2, leading to the opposite outcome contingencies. Crucially, participants did not know the current block’s contingency parameter and were required to relearn the contingency during each block. Please note that participants’ perceptual choices did not affect the delivery of rewards.

Figure 1c shows the behavior of an example participant. The task state (State) that was unobservable for participants was either 0 or 1 and determined which Gabor patch had the higher/lower randomly sampled contrast (Contrast difference). Due to the randomly varying strengths of the contrast difference, participants were confronted with perceptual uncertainty about the task state. In response to the participants’ economic choices (Economic choice), the task delivered a reward of either 0 or 1 point. (Reward). Based on these rewards, participants could sequentially learn the contingency parameter. As a result of perceptual uncertainty, participants faced a credit-assignment problem regarding the contingency between the hidden task state revealed through the Gabor patches, fractals, and reward.

### Perceptual uncertainty reduces learning accuracy

We commence with descriptive behavioral analyses of participants’ perceptual and economic choices. Subsequently, we analyzed the data using a model-based approach aimed at a mechanistic understanding of learning under perceptual uncertainty. We defined perceptual-choice performance as the observed frequency of a perceptual decision of the true but unknown high-/low-contrast Gabor patch. The mean ± standard error of the mean (SEM) perceptual-choice performance was 0.857 ± 0.008 (Figure 1d). The corresponding psychometric function shows that the accuracy of perceptual choices increased as a function of the contrast difference between the Gabor patches (Figure 1e, for details, see Psychometric function), and suggests that perceptual uncertainty was higher when contrast differences were smaller. We defined economic-choice performance as the observed frequency of an economic decision of the fractal with the higher reward probability. The mean ± SEM economic-choice performance was 0.78 ± 0.013 (Figure 1f). Participants’ economic-choice performance increased over the trials (Figure 1g), which indicates that despite perceptual uncertainty, they were able to learn the correct state-action-reward contingency throughout a block.

To test how well participants were able to learn the task in the absence of perceptual uncertainty, we also conducted an economic decision-making control experiment. Here, we did not sample the contrast differences at random but constantly presented Gabor patches with clearly different contrasts. In effect, this experiment corresponded to a conventional reward-based learning experiment without considerable influences of perceptual uncertainty. The perceptual-choice performance was 0.995 ± 0.001 (Figure 1d). To test if participants’ perceptual decision-making performance differed between the GB and control experiment, we conducted a two-sided paired sample t-test, which yielded a significant difference between the two conditions (*t* = 18.487*, p* < 0.001). Together, these results suggest that participants suffered from considerable perceptual uncertainty in the GB experiment and had low or nil perceptual uncertainty in the control task.

Furthermore, the mean ± SEM economic decision-making performance in the control task was 0.904 ± 0.011 (Figure 1f). As for the GB experiment, performance increased as a function of trials (Figure 1g). Finally, to test if economic decision-making performance under varying perceptual uncertainty levels in the GB experiment differed from performance in the control experiment without perceptual uncertainty, we used a two-sided paired sample t-test. This test yielded a significant mean difference between the conditions (*t* = 11.21*, p* < 0.001), and suggests that the additional perceptual uncertainty in the GB experiment considerably increased the difficulty to learn the reward contingency. For all choice performances plotted separately for each participant, we refer to S2 Single participant performance.

### Bayesian-inference and Q-learning agents

We designed several agent-based computational models to study the computational mechanisms of the learning dynamics (Figure 2a). In all agents, we assumed that perceptual uncertainty emerges because they suffer from perceptual noise in their visual system (Bach and Dolan, 2012). During the observation of the Gabor patches, the agents were not able to directly observe the presented contrast differences (Figure 2b, upper panels) but observed a noisy percept that corresponded to the actual contrast difference under the addition of Gaussian noise (Figure 2b, middle panels). To model this form of internal perceptual uncertainty, we applied a standard Bayesian binary hypothesis testing scheme (Levy, 2008) for computing the probability of the task state *s*_t_ conditional on the noisy observation of the contrast difference *o*_*t*_, 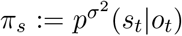. Here, the *σ* parameter indicates the agent’s level of perceptual sensitivity during the observation of the Gabor-patch contrast differences. That way, we obtained the agent’s belief over the states (belief state). High levels of perceptual sensitivity are associated with a less noisy perception of the Gabor patches and more distinct belief states (Figure 2b, left column). In contrast, lower perceptual sensitivity is associated with a more noisy perception of the patches and less distinct belief states (Figure 2b, right column). During perceptual decision making, the agents then selected the Gabor patch with the subjectively higher belief state. For more details about the agents’ perceptual-inference and choice mechanisms, see GB agent models.

**Figure 2.**
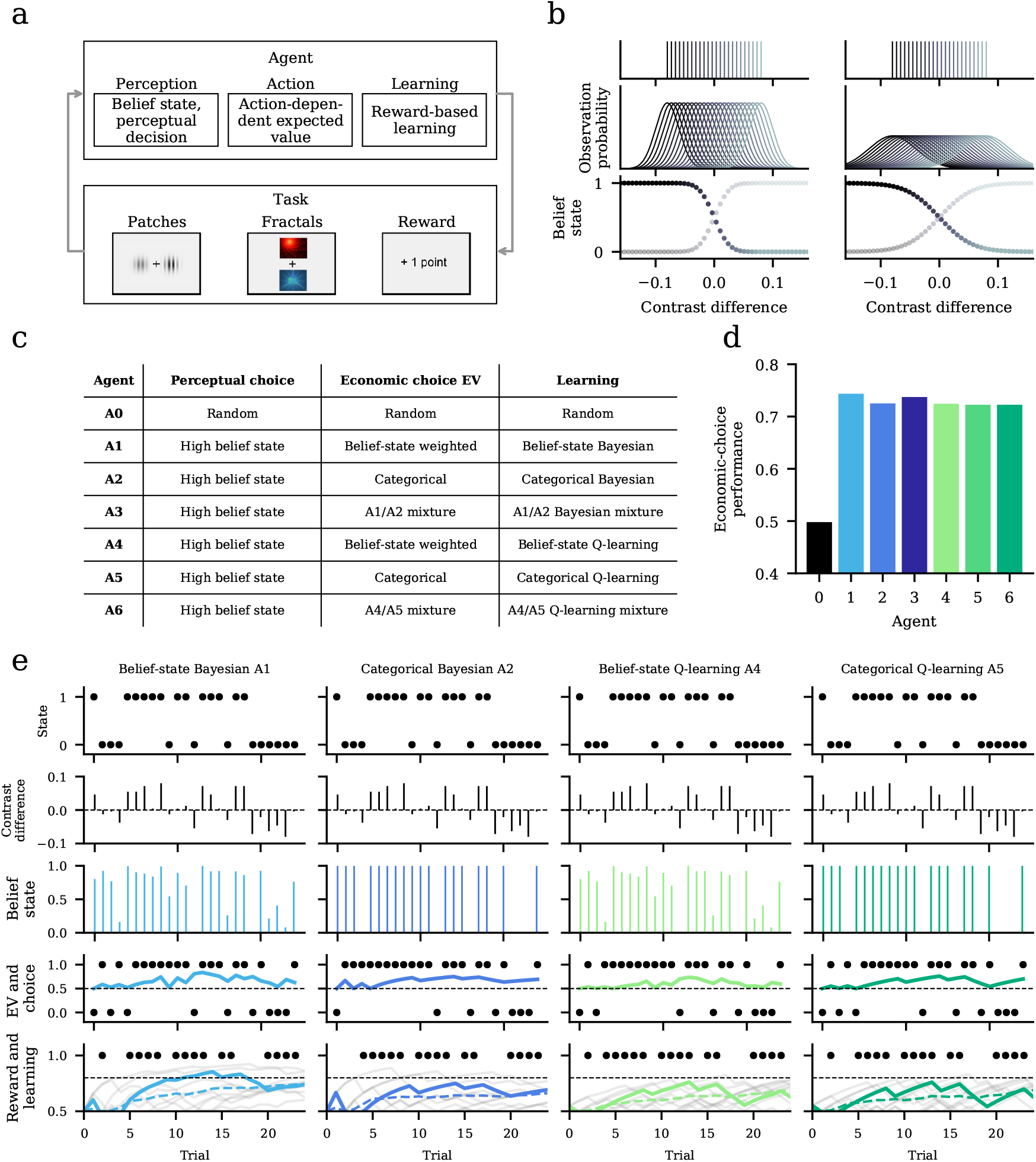
Agent-based computational modeling framework. **a)** The computational mechanisms of the agent-based models can be divided into a perception-, action-, and learning stage. In the perception stage, the agents perceived the contrast difference of the Gabor patches, computed the belief state expressing the current task state’s probability, and indicated their perceptual choice. In the action stage, the agents computed the expected value of the choice options under the consideration of the belief state and subsequently made their economic decisions. In the learning stage, the agents updated the inferred contingency parameter based on the received reward and belief state. **b)** We assumed that presented contrast differences (upper panels) were translated into noisy observations (Observation probability). The noisiness of the observations depended on the level of perceptual sensitivity. The left plots show an agent with intermediate perceptual sensitivity (sensory sensitivity parameter *σ* = 0.02), the right plots show an agent with low sensitivity (*σ* = 0.04). The lower panels illustrate the relation between observations and belief states, which reflect the observation-dependent probability of being in *st* = 0 or *st* = 1. The agent with higher perceptual sensitivity has more distinct belief states. **c)** Overview of the perceptual-choice, economic-choice, and learning properties of all agent models. Except for the random choice model A0, all agents chose the high belief state (BS) Gabor patch. However, the agents differed in how they computed expected values (EV) during economic decision making and how they learned the contingency parameter. In the economic-choice stage, expected values reflect the fractals’ expected reward under consideration of the belief state. The learned contingency parameter reflects the state-action-reward contingency of the current block (e.g., in state 0, reward probability of red fractal 80% (blue fractal 20%) and vice-versa for state 1). **d)** Simulated economic-choice performance of the agent-based models assuming low perceptual sensitivity (*σ* = 0.04) and deterministic economic-choice behavior based on the same task data as in the Gabor-bandit experiment. **e)** Task-agent interaction illustration. In the last subplot of each column (“Reward and learning“), the solid line shows the learned contingency parameter of the depicted block; gray lines represent ten example simulations, and the dashed line shows the average across simulations.

We assumed different reward-based learning and economic decision-making mechanisms across the agents (Figure 2c). The Bayesian belief-state agent (Figure 2e) explicitly considered the belief state during economic decision making and learning. During economic decision making, the agent combined the belief state and the learned contingency parameter to compute the fractals’ expected value. The expected value decreased when belief states were more similar (i.e., belief states close to *π*_*s*_ = (0.5, 0.5)). In the case of maximal perceptual uncertainty, the fractals’ expected value was (0.5, 0.5). Crucially, during learning, the agent utilized the belief state to weight learning of the contingency parameter optimally. That way, updates of the learned contingency were lower under high perceptual uncertainty. In the case of maximal perceptual uncertainty, the agent did not update the belief about the contingency parameter at all (see Figure SM 3). In brief, the belief-state Bayesian agent showed more cautious learning behavior under perceptual uncertainty to avoid that misinterpreted perceptual information corrupts learning.

The categorical Bayesian agent (Figure 2e) computed categorical belief states for learning and economic decision making that were determined by the agent’s perceptual decision (compare belief-state subplot to agent A1). After the agent chose the left Gabor patch, its belief state was *π*_*s*_ = (1, 0) and vice versa if the right patch was chosen. Consequently, during economic decision making, the fractals’ expected values were directly dependent on the current perceptual decision and not weighted as a function of perceptual uncertainty over the task state. Similarly, reward-contingency learning was determined by the current perceptual decision and unaffected by the agent’s perceptual uncertainty. The Bayesian mixture model (A3) combined the Bayesian belief-state and categorical agent. The free parameter *λ* determined the weight between the two agents, which allowed us to model a mixed consideration of belief states and categorical influences of the previous perceptual choice.

The belief-state Q-learning agent (Figure 2e) used the belief state to weight its expected values during economic decision making and its learning rate during reward-based learning of the reward contingencies. Moreover, the categorical Q-learning agent ignored perceptual uncertainty, i.e., represented a categorical belief state determined by the perceptual choice and not by uncertainty over the task state. In the Q-learning mixture model (A6), both belief states and categoricalchoice biases determined the agent’s behavior similar to the above. Finally, we used a control model (A0) that missed a mechanism for state inference and reward-based learning and, therefore, randomly generated perceptual and economic decisions. See GB agent models for details.

To compare the agent-based computational models, we conducted a simulation study independent of the participants’ choice behavior. We used a low level of perceptual sensitivity (σ = 0:04); that is, the agents suffered from relatively high perceptual uncertainty. Moreover, we assumed the absence of decision noise (softmax inverse temperature parameter of *β* = 100) to study purely exploitative economic choices. For the Q-learning agents A4-A6, we used a low learning-rate parameter (*α* = 0:1), and finally, for the mixture models A3 and A6, we used a mixture parameter leading to an equal weight of both belief states and categorical-choice biases (λ = 0:5). All agents except the random choice model A0 reached a considerable economic-choice performance (approximately 70-75%; Figure 2d). However, despite these comparable performance levels, the agents differed considerably concerning the accuracy with which they learned the task’s reward contingency. As a consequence of an optimal consideration of the belief state, the belief-state Bayesian agent (A1) learned the true but unknown contingency parameter of 0.8 accurately. In contrast, all other agents showed a tendency to underestimate the reward contingencies (see Figure 2e last row and Figure SM 5 for more details).

### Belief states and categorical-choice biases determine learning

To test which agent model explains our behavioral data most accurately, we developed a GB task-agent data-analysis model that allowed for estimating the seven above-described agent models (see GB task-agent data-analysis model and Parameter estimation).

As a measure of the agent models’ face validity, we compared the average performance achieved by human participants and simulated task-agent interactions based on the individually estimated participant parameters (Figure 3a). For the simulations, we took the same states and contrast differences of the blocks that our participants completed and repeated this procedure ten times to obtain a smoother average over multiple simulations. These simulations suggested that the Bayesian (A3) and Q-learning (A6) mixture models where belief states and categorical-choice biases controlled learning, may capture the data well. The economic-choice performances of both models were qualitatively close to participants’ data. Moreover, the simulation results suggested that the Bayesian belief-state agent that normatively considered perceptual uncertainty (A1) and the belief-state weighted Q-learning agent (A4) could qualitatively account for the choice data. Finally, the analysis indicated that purely categorical learning and decision-making strategies deviated from the data. Thus, both the categorical Bayesian (A2) and Q-learning agents (A5) achieved considerably lower performance than participants.

**Figure 3.**
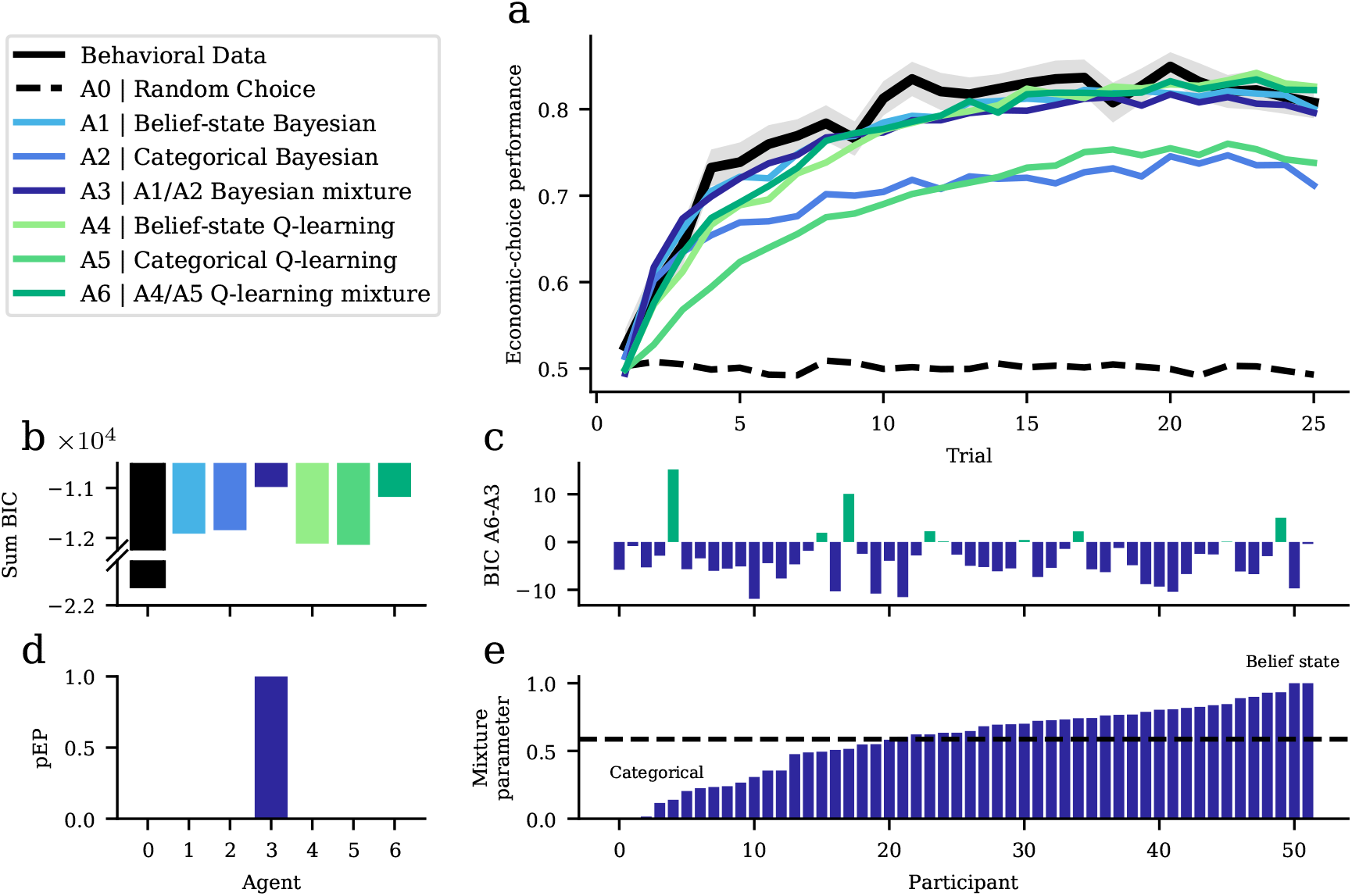
Agent model-based analyses. We analyzed participants’ behavioral data using the agent-based computational models. **a)** Based on several freely estimated parameters of the agent models, we simulated the models’ economic-choice performance. Compared to the empirical data, the normative belief-state Bayesian agent A1, Bayesian mixture model A3, belief-state Q learning model A4, and Q-learning mixture model A6 show qualitatively similar learning curves. In contrast, the categorical Bayesian agent A2 and categorical Q-learning agent A5 strongly deviate from the empirical data. **b)** Cumulated Bayesian information criterion (BIC) scores over participants of each agent model. Higher values indicate a better fit of the agent models to the behavioral data, revealing that the Bayesian mixture model A3 provides the best description of participants’ choice data. **c)** BIC difference between the Q-learning mixture model A6 and Bayesian mixture model A3. Negative values indicate a better fit of Bayesian mixture model A3, i.e., in 43 out of 52 participants, the BIC favored A3 over A6. **d)** Protected exceedance probabilities (pEP) similarly favor agent model A3. Together, b-d thus indicate that both belief states and categorical-choice biases determined learning and economic decision making in our participants. **e)** We next examined individual differences in the contribution of these factors according to the estimated mixture parameter of the Bayesian mixture model A3. A parameter of 1 indicates a full reliance on belief states, that is, normative belief-state Bayesian learning as in agent A1. In contrast, 0 indicates the complete dependency on the categorical-choice bias (i.e., no consideration of normative belief states as in the categorical Bayesian agent A2). Thus, in most participants, learning and economic decision making under perceptual uncertainty were determined by both belief states and categorical-choice biases. The dashed line indicates the mean parameter value (*λ* = 0.582).

To formally compare the agent models in light of participants’ choice data, we evaluated the cumulative BIC scores over participants for each agent model (Figure 3b). These indicated that the Bayesian mixture model (A3) explained the behavioral data best, followed by the Q-learning mixture model (A6). Next, we compared the BIC scores between these two models in more detail (Figure 3c), which indicated that in most participants (43/52) the Bayesian agent described the behavioral data better than the Q-learning agent. Moreover, assessing model plausibility using a random-effects Bayesian model selection procedure confirmed this result by allocating a protected model exceedance probability of more than 0.99 to the Bayesian mixture model (Figure 3d). Together, these results thus indicate that both belief states and categorical-choice biases modulated reward-guided learning under perceptual uncertainty in our participants.

To illustrate to which degree our participants relied on the belief state versus their categorical choices during learning and economic decision making, we extracted the mixture parameters where zero indicates a purely categorical strategy (i.e., categorical Bayesian learning) and one a full consideration of the belief state (i.e., belief-state Bayesian learning). Most participants showed a mixture between the two strategies, and the average *λ* parameter was 0.582 (Figure 3e). Next, to indicate to which extent the belief-state Bayesian agent described the data better than the random-choice control model A0, we computed a *pseudo-r*^2^ measure expressing the normalized difference between the negative log-likelihoods of agent A3 and A0 (Camerer and Ho, 1999; Daw, 2011). This revealed a *pseudo-r*^2^ of 0.475 and suggested that our winning model captured the choice data well overall. We additionally computed that the average probability with which the model successfully predicted participants’ choices was 0.7, which similarly indicates that the model explains a considerable amount of the variance between choices (for details see *Pseudo-r*^2^ and prediction accuracy).

Finally, we asked why the Bayesian mixture model captured the data better than the Q-learning mixture model. To this end, we also computed the *pseudo-r*^2^ of the Q-learning agent (*pseudo-r*^2^ = 0.47) and compared this measure to the winning model, which revealed that the *pseudo-r*^2^ of agent A6 was only minimally lower compared to the Bayesian belief-state agent (*pseudo-r*^2^ difference 0.005 in favor of agent A3). Along the same lines, the average probability with which the Q-learning agent predicted participants’ choices was the same as the predictive accuracy of the Bayesian agent (0.7). Together, these results indicate that the average ability to predict participants’ choices of the two models is similar. This suggests that the Bayesian mixture model wins the model comparison because it is more parsimonious than the Q-learning mixture model, which additionally uses a free learning-rate parameter that is penalized in the BIC-based model comparison. In both agents, the combination of belief states and categorical-choice biases seems to be the crucial aspect that leads to a better fit to the data than the other models in the model space. In summary, this result supports the conclusion that our participants used belief states to regulate learning and decision making under perceptual uncertainty, albeit with categorical-choice biases that distort a full consideration of belief states.

## Discussion

In natural settings, learning is often plagued by perceptual uncertainty because choice options or obtained outcomes can only partially be observed, which impairs the correct credit assignment between them. The critical problem in such environments is that in the presence of uncertain perceptual and reward information, an unexpected outcome can be the consequence of variability across rewards (e.g., a berry tastes bad because of natural variability) or misinterpreted perceptual information (two varieties of berries look similar and the wrong one has been eaten). While previous work in humans and other animals has extensively studied reward-based learning in situations with clear perceptual information (Dreher and Tremblay, 2016; Glimcher and Fehr, 2013; Rangel et al., 2008; Rushworth and Behrens, 2008), it remained unclear how humans learn under perceptual uncertainty. Therefore, this work aimed to examine to which degree humans regulate learning according to perceptual uncertainty and categorical-choice biases by applying several Bayesian-inference and Q-learning models.

First, we have shown how perceptual uncertainty should regulate learning from probabilistic reward feedback. In a normative Bayes-optimal learning model, we used a belief-state representation to formalize perceptual uncertainty (Daw, 2014; Dayan and Daw, 2008), which indicated that more uncertain belief states should render learning more cautious. In particular, when perceptual information is maximally uncertain, feedback should be entirely ignored. This Bayes-optimal agent could be dissociated from a Q-learning agent that scaled the learning rate according to the belief state (Chrisman, 1992) (see Figure SM 5). Second, we have shown that human participants utilize perceptual uncertainty to regulate learning and economic decision making but also identified a robust categorical-choice bias (Figure 3e). Our findings highlight that a perceptual choice that is reported before an economic choice and the reception of reward feedback nudges learning in the direction of this perceptual choice. In particular, perceptual choices render belief states more categorical, i.e., subjectively more confident, which leads humans to update reward contingencies after feedback more strongly compared to an optimal model. Finally, our model comparison indicated that a Bayesian agent that captured this categorical-choice bias provided a better description of the data than a Q-learning agent with a similar bias. This difference was primarily a consequence of the Q-learning model’s higher complexity because it contained an additional free learning-rate parameter. The finding that both the Bayesian and Q-learning model with categorical-choice bias described the data better than models without this bias reinforces the conclusion that both belief states and categorical-choice biases determine reward-guided learning under perceptual uncertainty.

Our Bayes-optimal agent model formally demonstrates the normative computations required to regulate learning under perceptual uncertainty effectively. The idea that belief states expressing the probability of the true but unknown task states guide learning under uncertainty is consistent with previous work (Babayan et al., 2018; Lak et al., 2017, 2020; Starkweather et al., 2017). However, in contrast to the current study, these previous studies combined belief-state inference with reinforcement learning models. Given that previous normative Bayesian models have successfully been used to model learning under various forms of uncertainty (Behrens et al., 2007; Mathys et al., 2014; Nassar et al., 2010; Payzan-LeNestour et al., 2013; Yu and Dayan, 2005), we were interested in whether optimal Bayesian-inference and Q-learning agents make dissociable predictions about human learning behavior. Therefore, we directly compared both types of models, which showed that although the belief-state Q-learning model scaled the learning rate according to perceptual uncertainty, the Bayes-optimal agent learned the reward contingency more accurately. At least concerning our model space and inference problem, these results indicate that a Bayes-optimal agent makes more efficient use of perceptual uncertainty and uncertain reward feedback to learn the reward contingency.

Importantly, the addition of the categorical-choice bias to both the belief-state Bayesian-inference and Q-learning agent was necessary to capture the participants’ choice data. That is, the confined but critical theoretical differences between Bayesian and Q-learning mentioned in the previous paragraph are not reflected in our empirical results, most likely as a consequence of the pervasive categorical-choice bias. This finding is consistent with previous studies in perceptual decision making, suggesting that perceptual uncertainty is not always optimally considered in a perceptual choice because of a categorical commitment to one particular interpretation of a stimulus (Fleming et al., 2013). Similar to our study, several studies have shown that past perceptual choices can induce such biases (Luu and Stocker, 2018; Stocker and Simoncelli, 2007; Urai et al., 2019). However, our results extend these studies by indicating that human participants consider prior categorical judgments about the most likely state of the environment during learning under perceptual uncertainty, which results in more substantial learning from reward feedback than dictated by a normative agent model.

One potential interpretation of this effect is that participants behaved as if the perceptual-, economic-choice, and learning stage were causally dependent (Luu and Stocker, 2018; Stocker and Simoncelli, 2007). In effect, learning and economic decision making were constrained by the perceptual choice and resulted in self-consistent behavior within a task trial. Luu and Stocker (2018) argued that self-consistency during perceptual inference might be the consequence of noise in the perceptual information’s working-memory representation. Thus, in our task, difficulties in maintaining an accurate representation of the contrast differences in working memory might have led participants to partly rely on the memory of the choice itself instead of exclusively on the observed contrast difference. Therefore, the belief state partly discarded decision-incongruent information in favor of self-consistency across the task stages (Peters et al., 2017). As argued in both Peters et al. (2017) and Luu and Stocker (2018), these seemingly sub-optimal behaviors might be resource-rational in natural settings with many hidden states and under limited cognitive capacities (see also Qiu et al. (2020)).

Our results are partly in line with previous work on reinforcement learning under perceptual uncertainty in the animal literature. Based on an experiment in monkeys, Lak et al. (2017) suggested that dopaminergic midbrain neurons compute reward prediction errors under consideration of belief states. In follow-up work in mice, Lak et al. (2020) additionally reported correlates of belief-state weighted expected values in the medial prefrontal cortex and found an effect of the previous trial’s belief state on current perceptual choices. Finally, in rats, Stolyarova et al. (2019) provided evidence for a broader system that combines perceptual and reward information in the service of learning, including the anterior cingulate cortex and basolateral amygdala. These studies raise the question, whether the identified brain areas similarly reflect belief-state weighted belief-updating mechanisms in humans. Conversely, our study raises the question of whether animals in these previous studies not only adjusted learning and decision making according to perceptual uncertainty but also according to the categorical-choice bias we identified. While the direct report of an isolated perceptual choice might have amplified this bias in our study, it is plausible that the brain categorizes perceptual information in the absence of an explicit choice report (Fleming et al., 2013), which would imply that categorical learning might still be at play in such tasks. Although different from our categorical-choice bias, the finding that the previous trial’s belief state affects current choices (Lak et al. (2020), see above) can be interpreted in this direction.

Future research could examine the neural representations of perceptual uncertainty and categorical-choice biases. Candidate areas involved in perceptual uncertainty computations from the perceptual decision-making literature are sensory, parietal, and frontal cortex (Kiani and Shadlen, 2009; Mulder et al., 2012; Summerfield and Koechlin, 2010; Van Bergen and Jehee, 2017; Van Bergen et al., 2015). There is some evidence to suggest that categorical biases emerging from the choice history are already reflected in early visual areas (Ester et al., 2020; Nienborg and Cumming, 2009). Moreover, Peters et al. (2017) showed that meta-cognitive confidence representations of choice-congruent evidence, which, as argued above, could be related to the categorical-choice bias, are distributed across multiple brain areas. Another interesting avenue for future research is to compare learning based on belief-state inference under perceptual uncertainty with other sources of uncertainty about task states. One research line shows that belief states modulate midbrain dopaminergic activity when animals learn under temporal state uncertainty (Starkweather et al., 2017) and uncertainty that results from different reward magnitudes (Babayan et al., 2018). Other studies implicate a broader system in reward-based learning based on hidden-state inference, including the orbitofrontal cortex (Schuck et al., 2016; Wilson et al., 2014). Together these studies could suggest that belief states arising from distinct sources of uncertainty might ultimately regulate learning in a common neural system. It is possible that categorical influences of past choices, as identified in our study and related factors from previous trials such as the stimulus history (Akrami et al., 2018), can also be identified as relevant in these sorts of learning problems.

One crucial issue that we have not addressed is dissociating the belief state’s influence on learning and economic decision making. In our models, we assumed that belief states similarly modulate expected values during economic decision making and reward-contingency learning after receiving feedback. Although our models accurately described participants’ choices, future research could directly ask participants to report their updated belief about the reward contingency in response to feedback (Nassar et al., 2010). A related extension is to ask participants to report their belief states, which might more completely capture the influence of our identified categorical-choice bias and other meta-cognitive influences beyond our model-based belief state measure (Fleming and Daw, 2017; Fleming and Lau, 2014; Peters et al., 2017; Stolyarova et al., 2019).

## Conclusion

To conclude, we examined human reward-based learning under perceptual uncertainty in a newly developed task that combined principles of two well-established tasks from perceptual and economic decision-making research. We first developed a Bayes-optimal agent model indicating that belief states should ideally lead to a more cautious regulation of reward-based learning under perceptual uncertainty to avoid that misinterpreted perceptual information corrupts learning. Second, we have shown that human participants consider belief states during reward-based learning and economic decision-making under perceptual uncertainty. However, prior perceptual decisions give rise to categorical-choice biases leading to a reduced consideration of perceptual uncertainty compared to optimal Bayesian inference. This categorical bias was the crucial factor of our various computational models that allowed us to capture human reward-based learning under perceptual uncertainty in both Bayesian and Q-learning agents. Taken together, our study highlights that humans dynamically combine uncertain perceptual and reward information during learning and decision making, yet categorical commitments to perceptually uncertain states of the environment modulate this integration substantially.

## Methods

### Participants

We recruited 54 participants from the local participant pool of Freie Universität Berlin and the Berlin School of Mind and Brain (Greiner, 2015). Participants provided written informed consent before partaking in the study. We excluded the data of two participants from all analyses due to a malfunction of the data-acquisition set-up. The effective study sample thus consisted of 52 participants. The participants of the effective study sample were between 18 and 33 years old (average age 24.4 years ± 0.6 SEM), 36 of them were women. Of the 52 participants, all reported normal or corrected-to-normal vision, one participant reported color blindness, and four reported currently remittent neurological or psychiatric diseases. The study was conducted in line with the human participant guidelines of the Declaration of Helsinki and was approved by the ethics committee of the Department of Education and Psychology at Freie Universität Berlin.

### Experimental procedure

We programmed the Gabor bandit (GB) task in PsychoPy2 (version v1.84.2) (Peirce, 2007). Before participants performed the GB experiment, they completed several training tasks as detailed in Supplementary Material S1.1 Participant training as well as both an economic and perceptual decision-making control task (S1.2 Control experiments). We conducted the experimental and training tasks on a Dell OptiPlex 780 computer (Dell Technologies Inc, USA) running Windows XP (Microsoft Cooperation, USA) in combination with a 19-inch SyncMaster 943BR monitor (Samsung Electronics Co., Ltd., South Korea). We asked participants to comfortably sit at a monitor distance of 60 cm and supported their heads by a chin-rest. For the detailed information that we provided about the relation between the relative Gabor-patch contrast locations and the reward probabilities of the fractal options, please see Supplementary Material S1.3 Participant instructions.

#### Trial structure

The GB experiment consisted of 12 blocks à 25 trials of a Gabor patch perceptual choice-augmented bandit task. Each trial of this task comprised three stages (Figure 1a): In the first stage, we simultaneously presented two Gabor patches (radius 5 cm, visual angle 9°) differing in their contrast levels at 6 cm to the left and right of a central fixation cross for 1 second. We asked 27 out of the 52 participants to choose the high-contrast Gabor patch and the remaining participants to indicate the low-contrast Gabor patch. Participants used the left and right cursor buttons of the computer keyboard and we allowed them to report their perceptual decision during the entire stimulus presentation time. We created the stimulus properties of the Gabor patches using the “GratingStim”^1^ function using a sine texture with a spatial frequency of 0.4 cycles per cm and a raised cosine alpha mask. We set the orientation and phase offset of the Gabor-patch stimuli to 0°. In addition, we manipulated the Gabor-patch contrast differences on a trial-by-trial basis to induce variable amounts of perceptual uncertainty about the current location of the high-/low-contrast patch. In particular, we controlled the contrast of a displayed patch *g* by manipulating the visibility *v*, which takes a value between 0 (equal to the background) and 1 (fully opaque). Accordingly, a presented Gabor patch thus corresponded to a weighted combination of the stimulus properties *z* (according to the texture, mask, orientation, and offset as described above) and the background color: *g* = *vz* + (1 − *v*)*h*. During all trials, we set the mean visibility of the patches to *v* = 0.5. To determine the relative contrast differences between patches, we used a uniform distribution of contrast differences extending over an interval of −0.08 (higher contrast on the left-hand side) to 0.08 (higher contrast on the right-hand side). Consequently, the absolute trial-by-trial visibility of the patches was in the range of *v* = 0.5 ± 0.04.

In the second stage of each trial, we displayed two clearly different, vertically aligned red and blue fractals for 1 second. We obtained the fractal stimuli from http://www.pixabay.com under a CC0 license and presented them with a width of 9.6 cm and a height of 6 cm and at a distance of 3 cm below and above the central fixation cross (visual angle 5° vertical and 9° horizontal). We asked participants to choose the fractal they thought was associated with the higher reward probability given the locations of the high- and low-contrast Gabor patches. Furthermore, we allowed participants to indicate their choice during the entire fractal presentation time window using up and down cursor buttons on a computer keyboard.

Finally, in the third stage, we presented a reward of zero (”+ 0 point” string) or one point (”+ 1 point” string) for 1 second. Between each trial stage, we presented a fixation cross for an inter-stimulus interval uniformly sampled from 0.9 to 1.1 seconds. Between trials, we presented a fixation cross for an inter-trial interval uniformly sampled from 0.9 to 1.1 seconds. During all stages, we presented the stimuli on a uniform gray background.

#### Task contingencies

The GB task’s central feature was the dependency of the fractal choice option reward probabilities on the relative locations of the high-/low-contrast Gabor patches, which were randomly assigned on each trial. For example, if the high-/low-contrast Gabor patch was displayed on the left side on a given trial, then the red fractal choice option was associated with a higher reward probability than the blue fractal choice option. In contrast, if the high-/low-contrast Gabor patch was displayed on the right side, then the red fractal choice option was associated with a lower reward probability than the blue fractal choice option (Figure 1b,c). Crucially, for 6 out of 12 randomly selected blocks, we reversed the state-action-reward contingency. Consequently, participants had to relearn the cumulative reward-maximizing contingency on each task block. In effect, throughout each block, participants faced a credit-assignment problem regarding the contingency of the high-/low-contrast Gabor-patch location (state), the fractal choice option (action), and the received rewards. For a detailed description of the probabilistic state-action-reward contingencies, please see GB task model. To exert experimental control over task statistics as experienced by the participants, we pre-generated the experimental sequence of trials on each block. In particular, we controlled for the distribution of contrast differences and rewards as well as the relative positions of the Gabor patches and choice fractals. For details, please see Supplementary Material S1.4 GB task statistics control. Finally, we repeated trials on which the participant failed to respond in either the perceptual or economic decision stage at the end of each block to avoid a preferential sampling of low perceptual uncertainty trials by the participants.

### Psychometric function

We computed the psychometric function (see Figure 1e) by binning the perceptual choices as a function of the presented contrast differences into 18 bins that spanned the range between −0.08 and 0.08. Subsequently, we computed the mean ± SEM across participants of the observed frequency of perceptual choices in favor of the right Gabor patch.

### GB task model

To render the GB task amenable to computational modeling, we first formulated a mathematical model of the task. In our documentation of this model, we follow the conventions of applied probabilistic modeling, i.e., we do not explicitly distinguish between probability distributions, probability density functions, or probability mass functions. Parameters are indicated by superscripts. We model a block of the task by the tuple

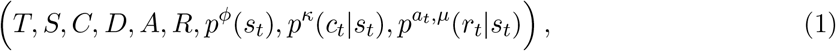

where

- *T* ≔ 25 denotes the number of trials per block, which are indexed as *t* = 1, 2*,…, T*,
- *S* ≔ {0, 1} is the set of task states governing the action-reward contingencies of the task,
- *C* ≔ [*−κ, κ*] with *κ* ≔ 0.08 is the set of Gabor-patch contrast differences,
- *D* ≔ {0, 1} is the set of perceptual choice options, where 0 and 1 represent the selection of the left and right Gabor patches, respectively,
- *A* ≔ {0, 1} is the set of economic choice options, where 0 and 1 represent the selection of the red and blue fractals, respectively,
- *R* ≔ {0, 1} is the set of rewards,
- *p^ϕ^*(*s*_*t*_) is the state distribution defined by the Bernoulli distribution

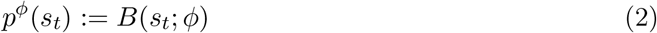

with state expectation parameter *ϕ* ≔ 0:5,
- *p*^*κ*^(*c*_*t*_|*s*_*t*_) is the state-conditional contrast-difference distribution defined by the product uniform distribution

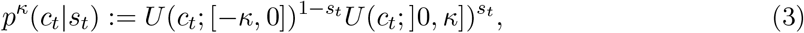

where contrast differences are thus negative for state 1 − *s*_*t*_ and positive for state *s*_*t*_, and
- 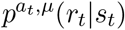 is the action- and parameter-dependent and state-conditional reward distribution representing the state-action-reward contingencies of the GB task. This distribution is defined as

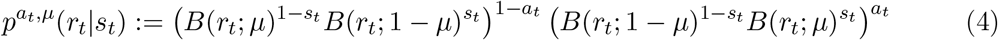

with contingency parameter *μ* ≔ 0.8 for half of the blocks and *μ* ≔ 0.2 for the other half of the blocks. Depending on the current state *s*_*t*_ and action *a*_*t*_, the task thus generates the rewards according to one of four possible Bernoulli distributions, two of which are identical.

In summary, on each trial, the task (1) samples a state *s*_*t*_ according to *p*(*s*_*t*_), (2) samples and displays a contrast difference *c_t_* according to *p^κ^*(*c_t_|s_t_*), (3) records a perceptual decision *d*_*t*_, (4) records an economic decision *a*_*t*_, and (5) samples and displays a reward according to 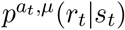.

### GB agent models

To formalize the putative cognitive processes of human participants interacting with the GB task, we next developed seven neuroscience-inspired agent models. These agents were of similar overall structure, that is, in all agents, we modeled perception of the Gabor patches and perceptual decision making and the computation of action-dependent expected values, economic decision making, and learning. However, the agents differed in their precise sequential-learning algorithms (see Figure 2a,c). In particular, we divided the agent-based computational models into a control model that exclusively generated random choices (A0), Bayesian-inference agents (A1-A3), and Q-learning agents (A4-A6). All artificial agents are implemented in *GbAgent.py* and *GbAgentVars.py* (see Software). In the following, we first introduce the perceptual-inference and -choice architecture common to all agents and then detail the economic-decision and learning components.

#### Perceptual inference and choice

For all agents, the perceptual model is represented by the tuple

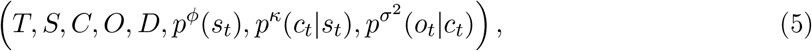

where

- *O* ∈ ℝ is a set of internal agent observations *o*_*t*_ that are assumed to result from the external Gabor-patch contrast difference *c*_*t*_ under additive perceptual noise,
- 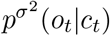 is the agent’s observation likelihood, defined as the contrast difference-conditional observation distribution (see Figure 2b)

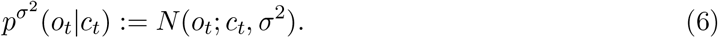

Crucially, the probability distributions of this tuple induce a joint probability distribution *p*(*s_t_, c_t_, o_t_*), allowing for evaluating an agent’s observation-conditional state distribution, modeling human perceptual inference, and for defining decision policies, modeling human perceptual decision making. In brief, each agent first evaluated its observation-conditional state distribution (belief state)

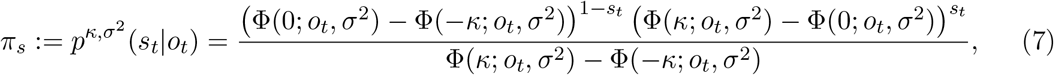

where Φ denotes the Gaussian cumulative distribution function (CDF). Subsequently, the agents chose the Gabor patch with the higher belief state, i.e., the patch with the subjectively higher/lower contrast

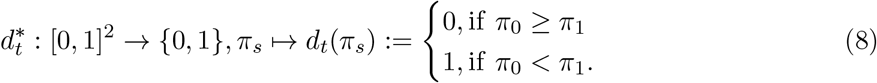

For mathematical details, please see Supplementary Material S3.1 Mathematical details of the perceptual model.

### Random choice model A0

#### Agent A0

Agent A0 was a random-choice model that lacked state- and state-action-reward contingency inference mechanisms. Because the belief state was always *π*_*s*_ = (0.5, 0.5), the agent was maximally uncertain about the current task state and generated random perceptual decisions. Furthermore, because agent A0 assumed a state-action-reward contingency parameter of *μ* = 0.5, it generated random economic choices.

### Bayesian inference agents A1, A2, and A3

All Bayesian inference agents are represented by the tuple

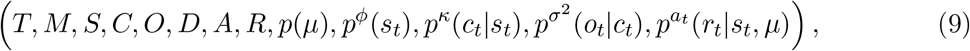

where

- in contrast to the task model, *μ* assumes the status of a random variable,
- *M* ≔ [0, 1] is the outcome space of this random variable,
- *p*(*μ*) is the agent’s task-block-specific initial uncertainty about *μ* and corresponds to a uniform distribution over *M*.

Here, the probability distributions of the tuple induce a joint probability distribution over *s*_*t*_, *c*_*t*_, *o*_*t*_, *d*_*t*_, *a*_*t*_ and *r*_*t*_. In addition to an analytical evaluation of an agent’s observation-conditional state distribution and the definition of perceptual decision policies as described above, this allows for formulating a sequential Bayesian state-action-reward contingency-parameter learning scheme to model human learning and memory. Moreover, this allows for defining decision policies in the fractal choice stage to model human economic decision making.

#### Agent A1

Agent A1 was a Bayes-normative exploitative model that explicitly considered its perceptual uncertainty during economic decision making and learning. We assumed that the agent made economic decisions such as to maximize its expected reward on each trial of the task, that is, for *t* = 1, 2*,…, T* selected 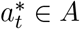 such that

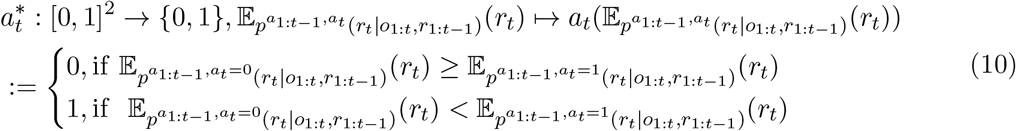

where the reward expectation conditional on action *a*_*t*_ = 0 (expected value) is given by

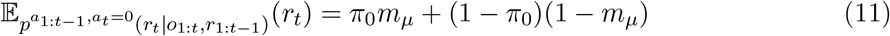

and the reward expectation conditional on action *a*_*t*_ = 1 is given by

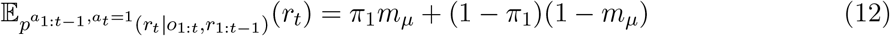

where

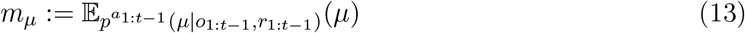

denotes the conditional expectation of the contingency parameter *μ* based on *a*_1:*t−*1_, *o*_1:*t−*1_ and *r*_1:*t−*1_. The action-dependent reward expectations thus conform to a convex combination of the expected probability for obtaining a reward, where the convex combination weighting parameters are given by the agent’s belief states *π*_0_ and *π*_1_, respectively.

In the learning stage, the agent updated the distribution of the state-action-reward contingency parameter

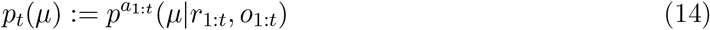

by considering the obtained rewards and the stimulus observations to weight the contingency-parameter update as a function of the current belief state. This distribution can be analytically evaluated as *t*th order polynomials in *μ* in a recursive fashion (Djurić and Huang, 2000). Specifically, the polynomial coefficients *ρ*_*t*,0_,…, *ρ_t,t_* ∈ ℝ of

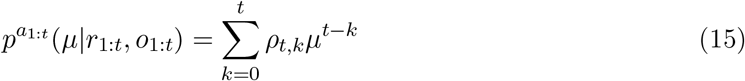

can be evaluated based on the coefficients *ρ_t−_*_1,0_*,…, ρ_t−_*_1*,t−*1_ of the polynomial

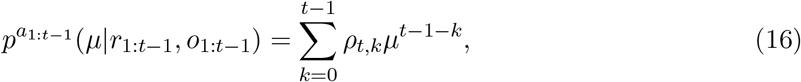

where

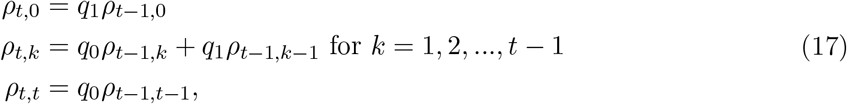

and where with

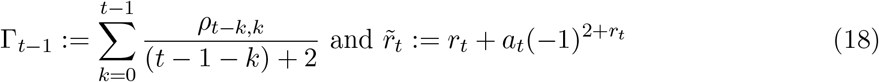

we have

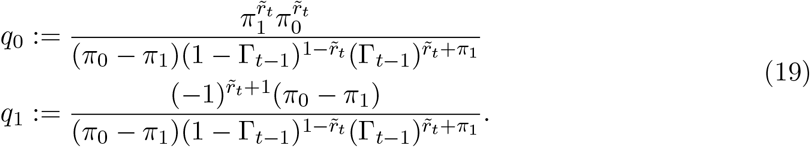

For mathematical details, please see Supplementary Material S3.2 Mathematical details of agent A1.

#### Agent A2

Agent A2 differed from A1 in that it ignored perceptual uncertainty during economic decision making and learning as a consequence of entirely relying on its perceptual decisions instead of on its stimulus observations. To model this categorical strategy, we adjusted the belief state to a binary state representation

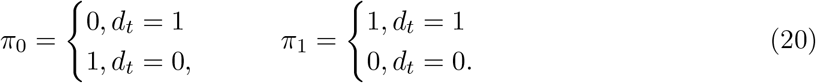

Thus, the belief state of agent A2 was entirely driven by the perceptual decision *d*_*t*_, and action-dependent reward expectations did not probabilistically factor in perceptual uncertainty. Similarly, during learning, the update of the state-action-reward contingency parameter was not modulated by perceptual uncertainty.

#### Agent A3

Agent A3 was a mixture model between agent A1 and A2, where the free parameter *λ* indicated the extent to which A3 behaved as agent A1 as opposed to agent A2. The action-dependent expected rewards were combined according to

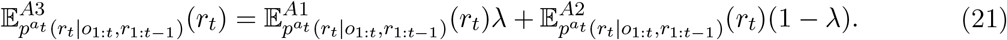

Similarly, the inferred contingency parameter was a linear combination of *p*_*t*_(*μ*) of agents A1 and A2, which was achieved by combining the polynomial coefficients of both models

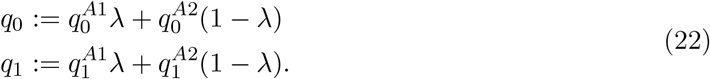

### Q-learning agents A4, A5, and A6

The Q-learning agents are represented by the tuple

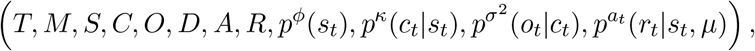

where in contrast to the Bayesian agent models, the distribution over the contingency parameter *p*(*μ*) is not included. Instead, the Q-learning agents sequentially updated state-action values. In particular, we considered a sequence of state-action-value functions

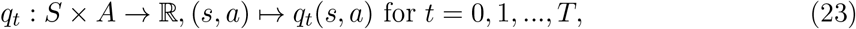

where *q*_*t*_(*s, a*) denotes the value that is assigned to state *s* and action *a* at trial *t*. At the beginning of each block, we initialized *q*_0_ for all state-action pairs to *q*_0_ = 0.5, reflecting the agent’s assumption that reward was equally likely for both actions.

#### Agent A4

Agent A4 considered the belief state during economic decision making and learning. This agent chose that action 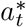, which maximized the probability to obtain a reward according to

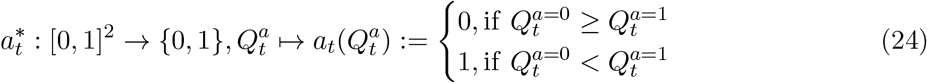

where

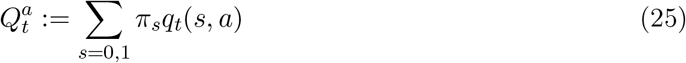

and where

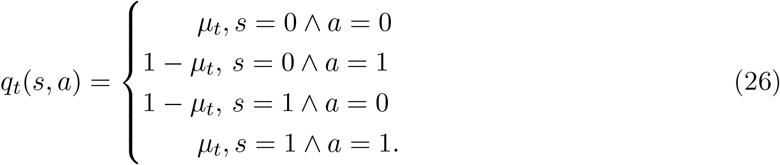

During learning, *μ*_*t*_ was updated under consideration of the belief state according to

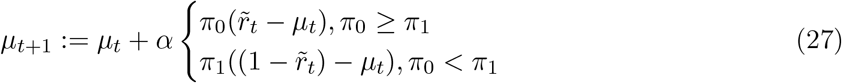

where 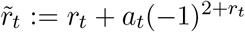 accounted for the action-dependency of the reward probability and
*α* was a free learning-rate parameter.

#### Agent A5

Agent A5 used a categorical economic decision-making and Q-learning strategy. That is, like agent A2, A5 represented a categorical belief state (eq. 20) and otherwise performed the same computations as agent A4.

#### Agent A6

Agent A6 was a mixture model between agents A4 and A5. During economic decision making, *Q*-values were combined according to

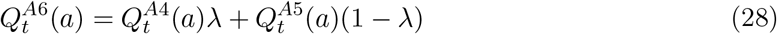

and during learning *q*_*t*+1_(*s* = 0*, a* = 0) according to

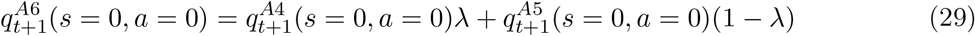

where *λ* was a free mixture parameter.

### GB task-agent data-analysis model

#### Perceptual decision making

In all agent-based models (except for the random choice model A0), the task-agent data-analysis model for the perceptual decisions corresponds to

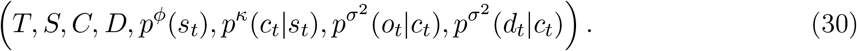

To take into account that we could not directly observe participants’ observations, we modeled perceptual decisions as random variables. In particular, we computed the probability of the perceptual decision *d*_*t*_ conditional on the presented contrast difference *c*_*t*_ as

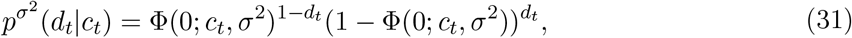

where Φ denotes the Gaussian CDF. For the corresponding proof, see Proof of (31).

#### Agent A1

Similarly, in agent A1 and the mixture component of A1 in the mixture model A3, the task-agent data-analysis model can be represented by the tuple

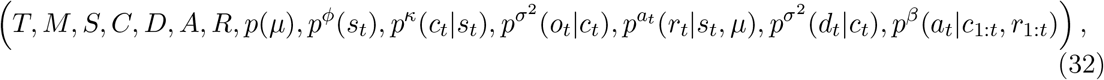

In this case, we modeled economic decisions as random variables based on two considerations. First, as in the case of perceptual decisions, we assumed that we could not access participants’ true observations. Given that the agent’s economic decisions were conditional on observations (cf. eqs. (10) - (13), we integrated over the range of contrast-difference conditional observations such that action-dependent reward expectations *v*_*a*_ were defined as

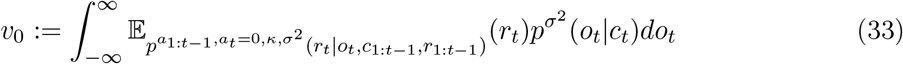

and

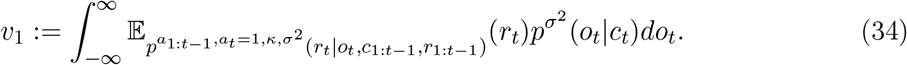

Second, as common in economic decision-making research, we assumed that action-dependent reward expectations were probabilistically translated into economic choices (Daw, 2011). Thus, we used the softmax choice rule to obtain the economic-choice probabilities that were conditional on the presented contrast differences and obtained rewards

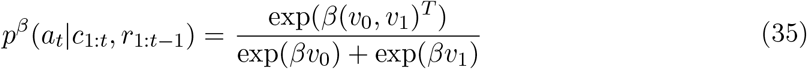

where the inverse temperature parameter *β* was estimated from the economic-choice data.

Finally, learning of the contingency parameter (cf. eq. (14)) also depended on the participants’ observations. To take into account that participants’ observations were unobservable for us, we integrated over the distribution of participants’ contrast-difference conditional observations when computing the posterior distribution over the contingency parameter:

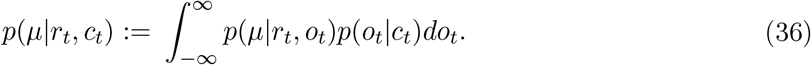

#### Agent A4

For the Q-learning agent A4 and the mixture component of A4 in the mixture model A6, the task-agent data-analysis model corresponded to the tuple

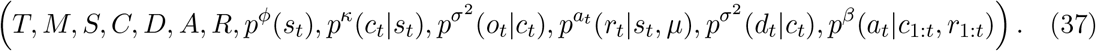

Similar to the above, during economic decision making we computed the action-dependent reward expectation (c.f. eqs. (24) - (26)) according to

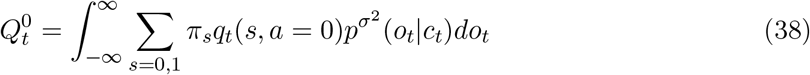

and

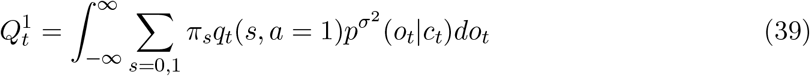

and obtained the corresponding choice probabilities by

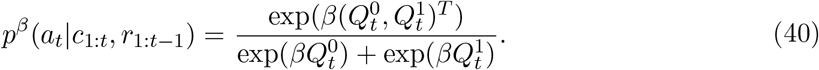

Finally, the learned contingency parameter *μ*_*t*+1_ of agent A4 in the corresponding task-agent data-analysis model (cf., eq. (27)) was computed as

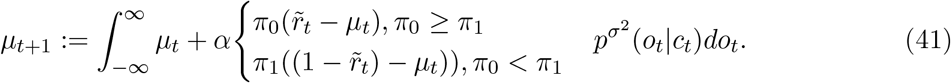

#### Parameter estimation

We estimated the free parameters with the bound-constrained optimization L-BFGS-B algorithm in Python 3.6 (Python Software Foundation; https://www.python.org/) via SciPy (Virtanen et al., 2020). The boundary constraints of the *σ* parameter were 0 and 0.1, of the *β* parameter 0 and 20, and of the *λ* and *α* parameters 0 and 1. For the corresponding code, please see *gb_estimation.py* in Software.

#### Model comparison

To compare the evaluated agent-based computational models, we first computed the Bayesian information criterion for each participant

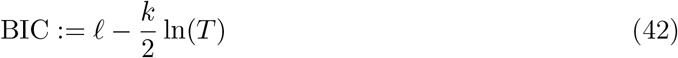

where next to the maximum log-likelihood *ℓ*, the number of free parameters *k*, scaled by the logarithm of the number of trials *T* is taken into account (Stephan et al., 2009). For model comparison on the group level, we subsequently computed protected exceedance probabilities, which indicated the probability that a particular model was more likely than any other model of the model space (Rigoux et al., 2014).

#### Model recovery

To test if we could distinguish the seven agent-based computational models, we conducted a model-recovery study (Figure SM 4a). With each model, we simulated data of *N* = 256 fictive participants where each data set consisted of 12 blocks á 25 trials, and where we used the states, contrast differences, and reward probabilities that participants dealt with in the GB experiment (for every simulation we randomly selected a data set of a participant). We used a large range of parameter values for the simulations. For agents A1-A6, the perceptual sensitivity parameter was *σ* ∈ {0.015, 0.025, 0.035, 0.045} and the softmax slope parameter *β* ∈ {4, 8, 12, 16}. For the mixture parameter of agents A3 and A6, we additionally used *λ* ∈ {0.2, 0.4, 0.6, 0.8} and for the learning-rate parameter of agent A4-A6, we used *α* ∈ {0.1, 0.3, 0.5, 0.7}. For each agent, we used all possible combinations of these parameters. Subsequently, we evaluated all agent models based on the simulated data and compared them using the BIC and protected exceedance probabilities. During model evaluation, we fixed all parameters to the true parameters that we used during the simulations.

#### Parameter recovery

To test if we could reliably estimate our free parameters, we conducted a parameter-recovery study under comparable task conditions as in our empirical experiments (Figure SM 4b-e). In particular, we submitted the simulated data of the 256 agents from the model-recovery study to the same model estimation analysis as for our empirical data. This analysis suggested that we could reliably recover the true parameter values in conditions similar to our experiments.

#### *Pseudo-r*^2^ and prediction accuracy

We computed a *pseudo-r*^2^ measure to express the fractional reduction in the negative log-likelihood of agent A3 (*ℓ^A^*^3^) toward *ℓ* = 0 (perfect prediction) relative to random predictions (agent A0; *ℓ^A^*^0^) (Camerer and Ho, 1999; Daw, 2011)

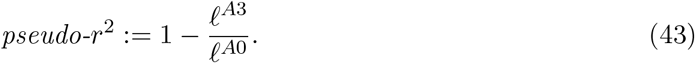

Moreover, we computed the average probability with which agent A3 predicted participants’ choices according to exp 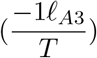, where *T* denotes the number of trials. Both steps were similarly applied to agent A6.

#### Software

The task was programmed in Psychopy2 (Peirce, 2007). The task code is available at https://github.com/rasmusbruckner/gaborbandit_task. Data analysis and computational modeling was performed in Python 3.6 (Python Software Foundation; https://www.python.org/). We used the NumPy (Oliphant, 2006), SciPy (Virtanen et al., 2020), pandas (McKinney, 2010), matplotlib (Hunter, 2007), seaborn (https://doi.org/10.5281/zenodo.12710) and tqdm (https://doi.org/10.5281/zenodo.1239851) libraries. Protected exceedance probabilities were computed in SPM12 (Penny et al., 2011) using Matlab 2017b (The Mathworks Inc., USA). Pseudonymized experimental data in brain imaging data structure (BIDS) format (Gorgolewski et al., 2016) and the Python analysis code are available at https://github.com/rasmusbruckner/gaborbandit_analysis.

## Acknowledgments

We thank Jan Bruckner, Petar M. Djurić, Vincent Boelens, Matt Nassar, Megan Peters, and Steve Fleming for helpful comments and discussions. During his Ph.D., R.B. was supported by the International Max Planck Research School LIFE, Berlin, Germany.

## CRediT author statement

Rasmus Bruckner: Conceptualization, Data curation, Formal analysis, Investigation, Methodology, Project administration, Resources, Software, Validation, Visualization, Writing – original draft, Writing – review & editing; Hauke R. Heekeren: Conceptualization, Funding acquisition, Project administration, Resources, Supervision, Writing – review & editing; Dirk Ostwald: Conceptualization, Methodology, Resources, Software

## Supplementary Material

### Data preprocessing

In the GB experiment, participants missed between 1 and 67 trials. The mean ± SEM number of misses was 10.99 ± 1.524. In the control experiment, participants missed between 0 and 9 trials. Here, the mean ± SEM was 2.846 ± 0.318. In both cases, missed trials were repeated at the end of the block. Consequently, each participant completed *T* = 300 in the GB experiment and *T* = 150 trials in the control experiment.

### S1. Additional experimental details

#### S1.1 Participant training

Before performing the GB and economic choice control experiment, participants performed a series of training tasks. Specifically, they first performed two blocks of a pure perceptual decision-making experiment that only involved the Gabor patches. The first training task comprised 50 trials with trial-wise feedback on the correctness of the perceptual decision, and the second comprised 100 trials without any feedback. Next, participants performed two training blocks à 25 trials of the economic decision-making control experiment with clearly distinct Gabor-patch contrasts (contrast differences −0.08 and 0.08). Finally, participants performed two training blocks à 25 trials of the GB experiment. The stimulus order of all training blocks was pre-generated as for the main experiment.

#### S1.2 Control experiments

The control experiment consisted of 6 blocks à 25 trials of the GB task for which the Gabor-patch contrast differences were fixed to either −0.08 or 0.08 and hence clearly perceptually distinguishable. Finally, before the main GB experiment, participants completed an additional perceptual-choice task that consisted of 100 choices between the high-/low-contrast Gabor patch (Figure SM 2a). Here, participants missed between 0 and 7 trials. The mean ± SEM number of missed trials was 2.577 ± 0.295. The perceptual-choice performance was similar to perceptual-choice performances in the GB experiment.

#### S1.3 Participant instructions

During the training phase, participants were instructed on the GB task as follows:

> *“You should now try to maximize your earnings on the task. There are two possible mappings between the position of the high-contrast image (left and right) and the two patterns (red or blue). The aim is to figure out which pattern you should choose, given the position of the high contrast image, to maximize the probability of a reward. You will learn this mapping from feedback after your choice. At the end of the task, you will be paid out the monetary equivalent of your collected points.”*

Moreover, we explicitly informed the participants about the two possible task states. Upon presentation of the task instructions above and in combination with an example trial, we advised the participants as follows:

> *“If the high-contrast patch is on the other side, you should also choose the other fractal. That is, if you prefer the red pattern given that the high-contrast image is on the left-hand side, then you should choose the blue fractal given that the high contrast-image is on the right-hand side.”*

The original German instructions corresponding to the two statements above read:

> *“Ab jetzt sollst Du versuchen, Deinen Gewinn in der Aufgabe zu maximieren: Es gibt zwei mögliche Kopplungen zwischen der Position des kontrastreichen Bildes (rechts und links) und den zwei Mustern (rot oder blau). Das Ziel ist es herauszufinden, gegeben der Position des kontrastreichen Bildes, welches Muster Du wählen solltest, um wahrscheinlich eine Belohnung zu erhalten. Diese Kopplung lernst Du durch Feedback nach der Antwort. Deine so verdienten Punkte bekommst Du am Ende, in Geld umgerechnet, ausgezahlt.”*

and

> *“Wenn sich das kontrastreiche Bild auf der anderen Seite befindet, wählst Du auch das andere Muster. Das heißt: Wenn Du bei “kontrastreiches Bild links” am besten das rote Muster wählst, so nimmst Du bei “kontrastreiches Bild rechts” am besten das blaue Muster.”*

#### S1.4 GB task statistics control

To match the experimentally presented task statistics with the underlying probabilistic properties of the task, we created the experimentally presented task blocks as follows. For each task block, we first sampled a trial sequence of 30 trials and then ensured that the statistics of these 30 trials matched the underlying task properties. Specifically, the 2nd to 26th trial of such a sequence was included for experimental presentation, if the sequence of 30 trials met the following criteria
 
- the average of absolute contrast differences over trials was in the range 0.036 to 0.044, which corresponds approximately to one standard deviation, and the standard deviation of absolute contrast differences over trials was between 0.021 and 0.025,
- the observed number of +1 and +0 rewards matched the contingency parameter of 0.8, i.e., the sequence contained 24 +1 rewards and 6 +0 rewards,
- on 15 of the 30 trials, the high-contrast Gabor patch was presented on the left.

In addition, we further increased the homogeneity of the experienced task statistics by only accepting the 2nd to 26th trial of the trial sequence for inclusion in the experimentally played-out task sequences, if

- on 15 of the 30 trials, the red choice fractal was presented above the fixation cross, and
- minimally 9 repeats of rewards or no rewards (1,1,1 or 0,0,0) and maximally 13 repeats to avoid extremely long/short sequences of the same outcome.

In Figure SM 1 below, we depict the average experimental task statistics across all task blocks and participants.

### S2 Single participant performance

In Figure SM 2 below, we plot the single participant perceptual- and economic-choice performances for all tasks.

#### S3.1 Mathematical details of the perceptual model

##### Belief state

As shown in Proof of (7) the agent’s observation-conditional distribution over task states at trial *t* can be derived in a straightforward manner from the marginal probabilistic model of interest

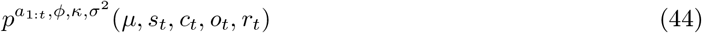

at trial *t* = 1*,…, T*, where Φ denotes the cumulative density function of the Gaussian distribution.

**Figure SM 1.**
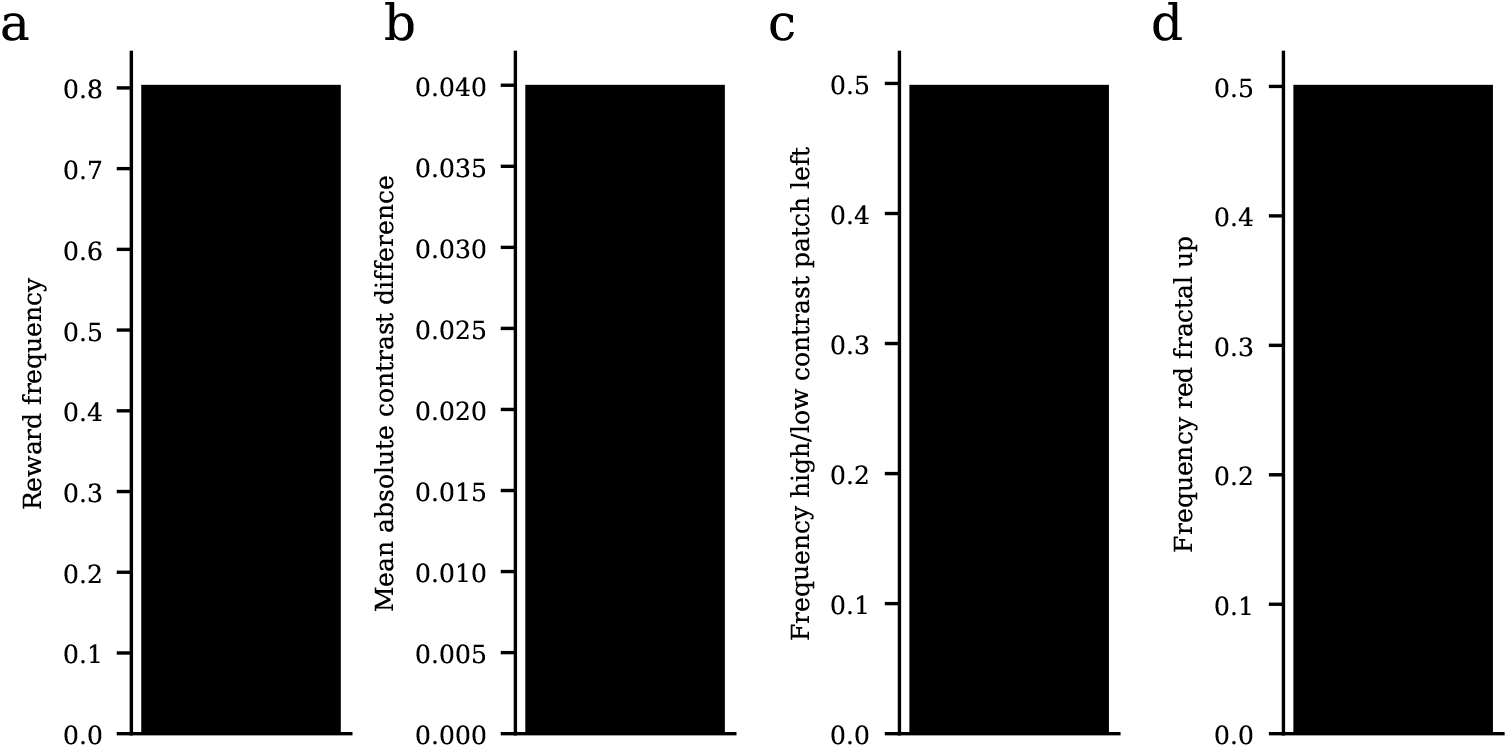
Experimental task statistics. **a)** Average reward frequency given high-reward economic choice. **b)** Mean absolute contrast difference. **c)** Frequency with which high-/low-contrast patch was presented on the left-hand side. **d)** Frequency with which red fractal was presented on the upper half of the screen.

**Figure SM 2.**
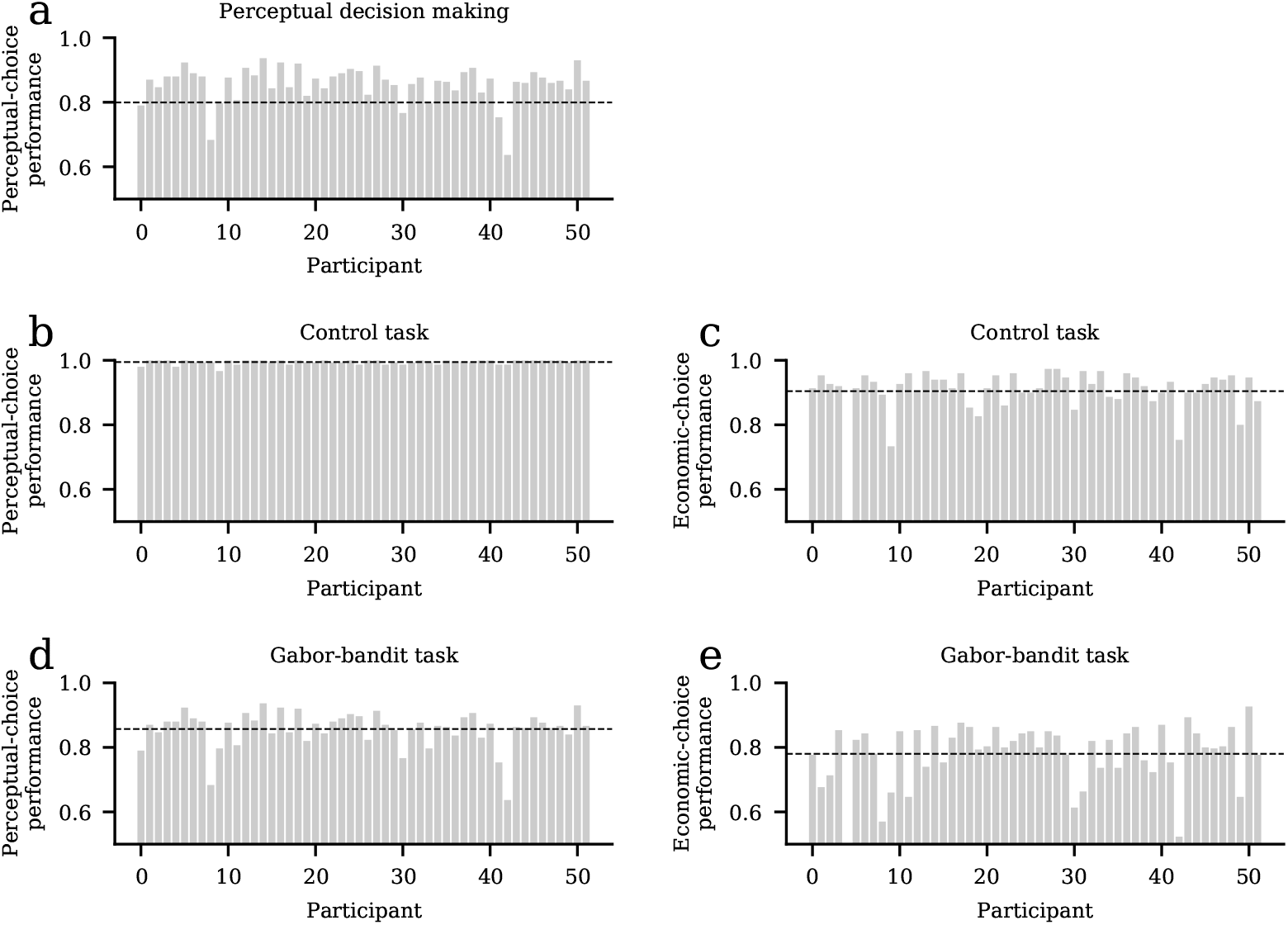
Single participant performance. **a)** Perceptual-choice performance in the perceptual decision-making control experiment. **b)** Perceptual-choice performance in the economic decision-making control experiment. **c)** Economic-choice performance in the economic decision-making control experiment. **d)** Perceptual-choice performance in the GB experiment. **e)** Economic-choice performance in the GB experiment.

##### Perceptual decision

The perceptual-choice policy shown in eq. (8) follows from the assumption that the agents aim at minimizing the 0-1 loss function

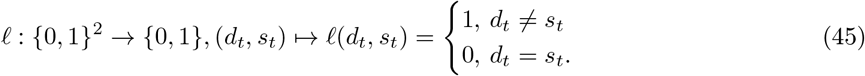

Eq. (8) then expresses the policy to choose that perceptual decision 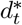 for which

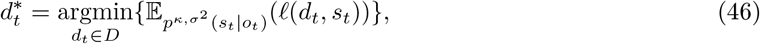

where 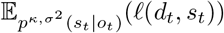 denotes the agent’s expected loss. Because the true state is unknown, the agent computes the expected loss using the the above-defined belief state that reflects the observation-conditional probability of the states:

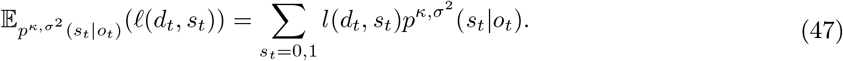

That is, for *d*_*t*_ = 0, the expected loss corresponds to

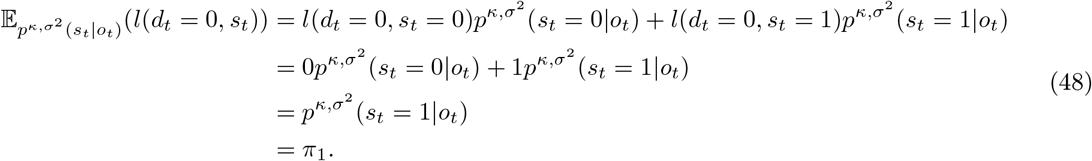

Likewise, for *d*_*t*_ = 1 we obtain

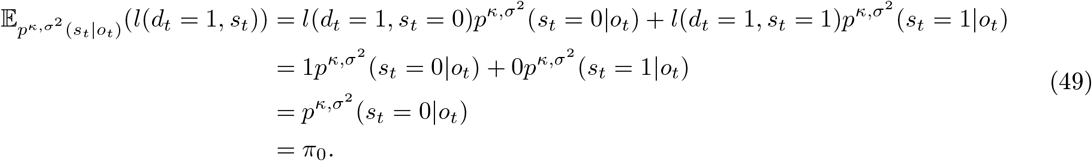

#### S3.2 Mathematical details of agent A1

##### Probabilistic agent representation

The definitions of the task and agent models in (1) and (9), respectively, induce a joint probability distribution over all random variables associated with the agent. This probability distribution is characterized by its conditional independence properties, which are given by

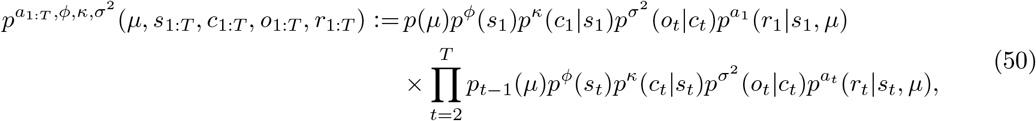

where

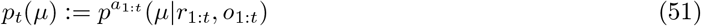

denotes the posterior distribution of *μ* on trial *t* which in a sequential Bayesian update acts as the prior distribution of *μ* on trial *t* + 1.

As discussed in Bayesian inference agents A1-A3, the agent selects, on each trial *t* = 1*,…, T*, the action 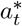 that promises the higher expected reward, given its observations of subjective contrast differences *o*_1:*t*_ and action *a*_1:*t−*1_-dependent rewards *r*_1:*t−*1_. Formally, the agent follows the policy

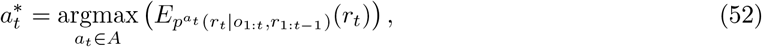

which requires evaluating the expectation of the random variable *r*_*t*_ under the conditional distribution 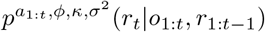 as a function of *a*_*t*_. This can be achieved by first showing that for the experimental setting *ϕ* ≔ 0.5, this conditional distribution can be written as

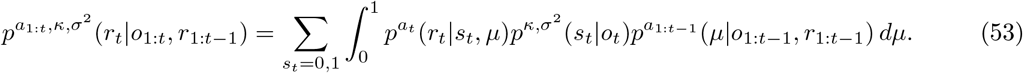

Eq. (53) is helpful, because explicit expressions are available for 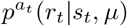 (by definition of the agent model), 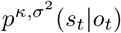 (as this corresponds to the agent’s belief state on trial *t*), and 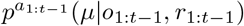 (as this corresponds to the contingency-parameter inference distribution on trial *t* 1). We prove (53) in Proof of (53). With the explicit form of (53), we then derive eqs. (11) and (12) in Proof of (11) and (12). In Proof of (14), we then proof the recursive forms of the polynomial coefficients of the agent’s contingency parameter probability density function. In Figure SM 3, we additionally illustrate the contingency parameter probability density function across several trials under no perceptual uncertainty conditions (Figure SM 3a), high perceptual uncertainty (Figure SM 3b), and maximal perceptual uncertainty (Figure SM 3c).

**Figure SM 3.**
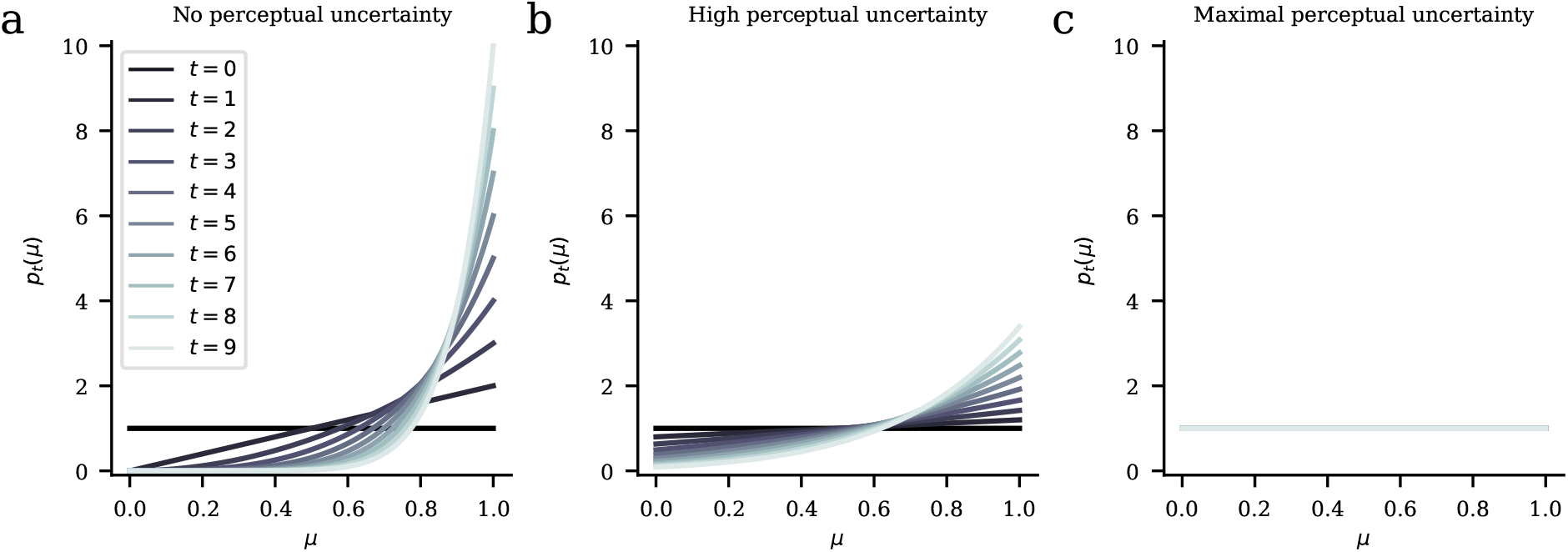
Agent contingency learning illustration. **a)** Evolution of the contingency parameter probability density function under conditions of no perceptual uncertainty (*π_s_* = [1, 0]), **b)** of high perceptual uncertainty *πs* = [0.6, 0.4], **c)** and maximal perceptual uncertainty (*π_s_* = [0.5, 0.5]).

##### Model- and parameter-recovery validation

To test which model explained our behavioral data most accurately, we developed a GB task-agent data-analysis approach that allowed for estimating the seven above-described agent models (see GB task-agent data-analysis model and Parameter estimation). In particular, we had no direct access to participants’ observations in the GB task because perceptual uncertainty arose from internal sensory noise in the participants’ perceptual (visual) system. Therefore, an estimation of the agent models’ parameters required an embedding of the agent models into a statistical framework that accounted for our uncertainty over participants’ observations (Daunizeau et al., 2010).

In a model-recovery analysis, we first tested if these agent models could be reliably dissociated from each other. Here we simulated experimental data with an agent model (e.g., agent A0), then evaluated the thus obtained data with each agent model (agent A0-A6) and subsequently tested which evaluated model described the data best. According to our Bayesian information criterion (BIC) (Stephan et al., 2009) and protected exceedance probability (Rigoux et al., 2014) based model comparison (see Model comparison), we were able to reliably dissociate all models from each other (Figure SM 4a). For example, when we simulated experimental data using agent A0, our model comparison indicated that agent A0 had the highest BIC and protected exceedance probability among the models in the model space. See Model recovery for details.

We also conducted a parameter-recovery study to test how well the individual parameters could be estimated. The random choice model A0 had no free parameters, and both perceptual and economic-choice probabilities were equal to (0.5, 0.5). For agent A1-A6, in order to determine the individual perceptual sensitivity of participants, we estimated the *σ* parameter, i.e., the standard deviation of the Gaussian distribution modeling perceptual sensitivity during the observation of the Gabor patches (Figure 2b). We estimated this parameter using perceptual choices, that is, the estimation was independent of the remaining free parameters to exclusively capture perceptual sensitivity during the perception of the Gabor patches (Figure SM 4b). Next, we estimated the additional free parameters based on economic choices to exclusively capture influences on economic decision making and learning. For the Bayes-optimal agent A1 and the categorical Bayesian agent A2, we estimated the slope parameter *β* of the softmax choice rule (Figure SM 4e). For the mixture model A3 we additionally estimated the *λ* parameter, determining the mixture weight between agent A1 and agent A2, where higher values indicated a higher contribution of agent A1 and thus a stronger consideration of the belief state (Figure SM 4c). Moreover, for the belief-state Q-learning agent A4 and the categorical Q-learner A5, we estimated the *β* parameter and the learning-rate parameter *α* (Figure SM 4d,e). Finally, in agent A6, we additionally estimated the *λ* parameter that determined the mixture between A4 and A5 (Figure SM 4c). Together, our parameter-recovery analysis suggests that we could reliably estimate the free parameters of all agent-based computational models.

**Figure SM 4.**
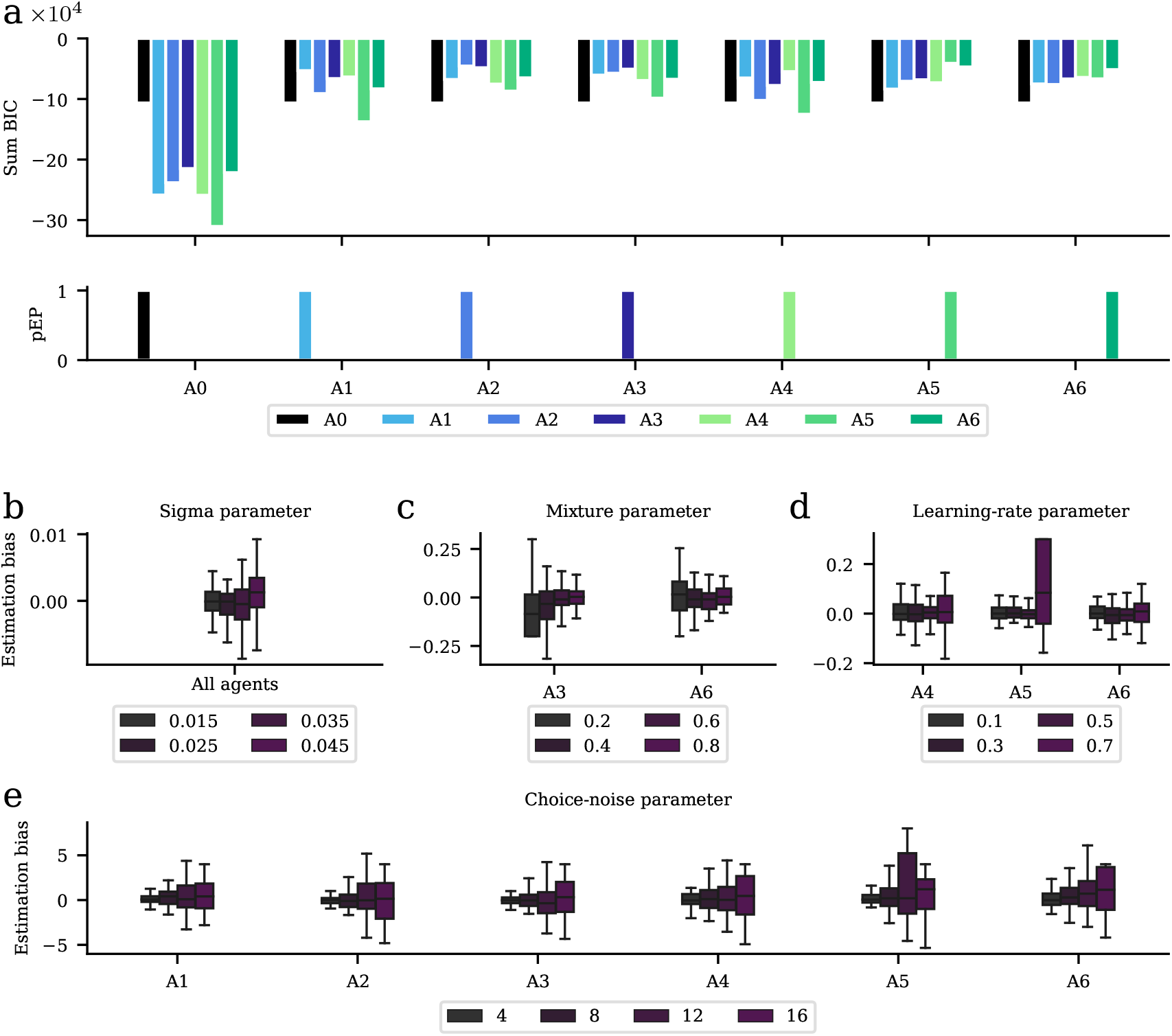
Model and parameter recovery. **a)** For our model-recovery analysis, we simulated data with each agent-based computational model using the same states and contrast differences as for our participants and a broad range of model parameters. We then evaluated the simulated data using each task-agent data-analysis model and compared the corresponding cumulated Bayesian information criterion (BIC) values and protected exceedance probabilities (pEP). Each agent model used to generate the data had larger BIC and pEP values than the other agent models that did not generate the data. This indicates that all models can reliably be recovered. b) In a parameter-recovery analysis, we submitted the simulated data to the same estimation pipeline as our empirical data and computed the difference between the estimated and generative parameters (estimation bias). This subplot shows the mean ± SD estimation bias of the sigma parameter that modeled perceptual sensitivity. c) Mean ± SD estimation bias of the *λ* parameter that modeled the mixture of the Bayesian (A1 and A2) and Q-learning agents (A4 and A5). **d)** Mean ± SD estimation bias of the learning-rate parameter *α*. **e)** Mean ± SD estimation bias of the *β* parameter that modeled economic-choice noise. Together, the parameter-recovery analysis indicates that all parameters can be reliably recovered.

**Figure SM 5.**
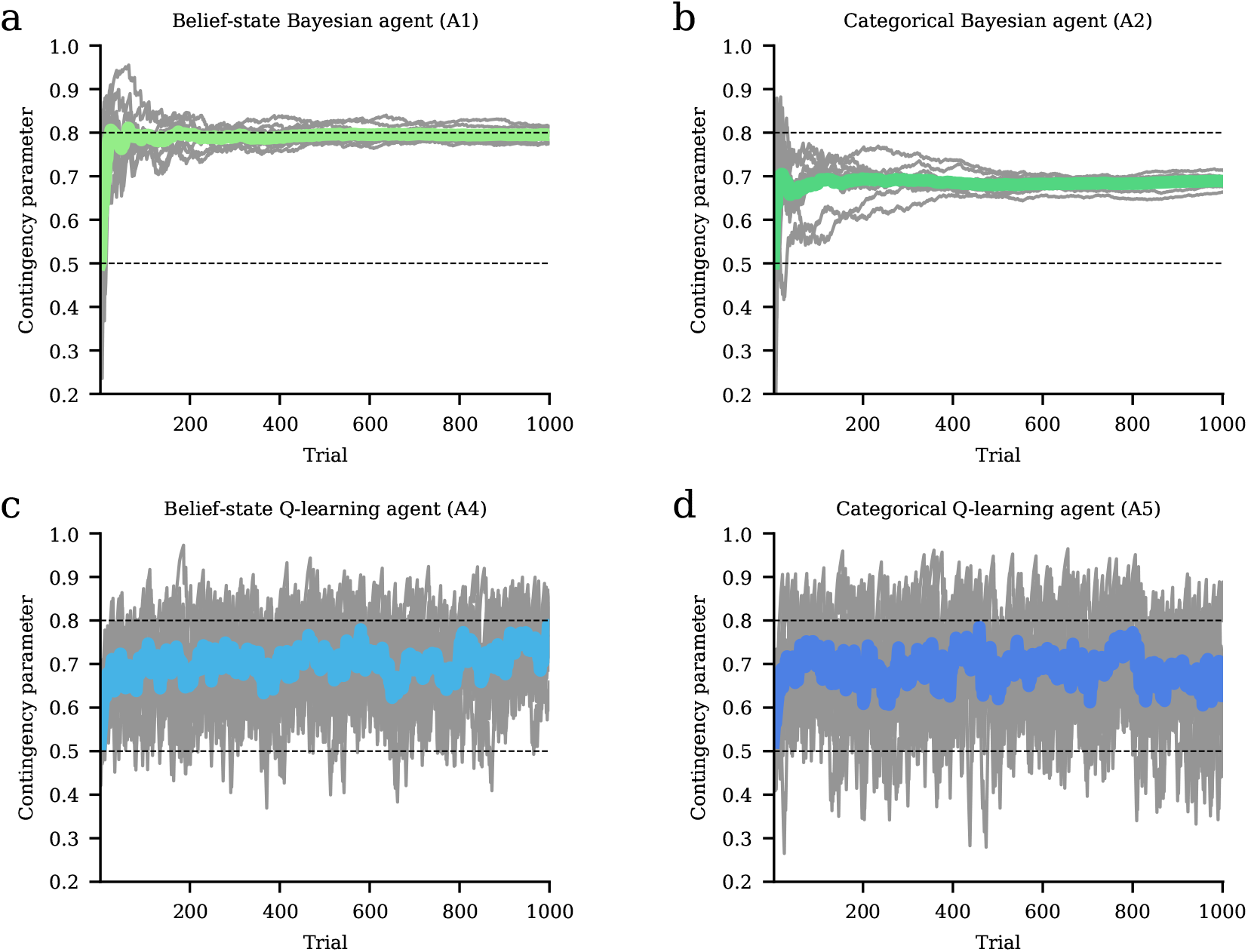
Learning accuracy of the agents. We compared the agents’ learned contingency parameters across 10 simulated blocks of 1000 trials. The true but unknown contingency parameter was equal to 0.8 (upper dashed line). **a)** This simulation indicated that the belief-state Bayesian agent (A1) accurately learned the contingency parameter. **b)** The categorical Bayesian agent showed an underestimation of the contingency parameter. **c,d)** For the belief-state and categorical Q-learning agents we used a learning-rate parameter of 0.1. Both agents underestimated the contingency parameter. In all agents, the perceptual-sensitivity parameter was *σ* = 0.04.

## 1. Supplementary Material Appendix

Notation. In the following, we denote the probability density function for the uniform distribution of a univariate random variable on the interval [*a, b*] ⊂ ℝ by

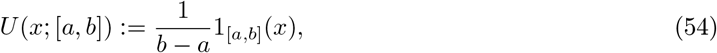

where 1_*A*_ denotes the indicator function for the set *A* ⊂ ℝ. We denote the probability mass function for the Bernoulli distribution of a random variable with outcome space {0, 1} by

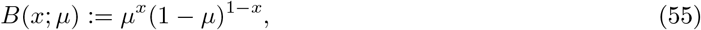

where *μ* ∈ [0, 1] denotes the expectation parameter. Finally, we denote the probability density function and the cumulative density function for the Gaussian distribution of a univariate random variable by

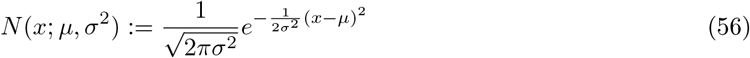

and

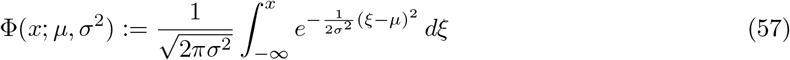

respectively, where *μ* ∈ ℝ and *σ*^2^ *>* 0 denote the expectation and variance parameters, respectively.

## 1.1 Proof of (7)

To show that (7) holds, we consider the marginal probabilistic model of interest at trial *t*

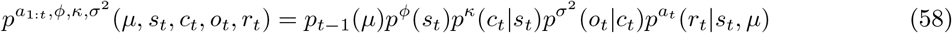

and directly derive the observation-conditional state distribution by evaluating

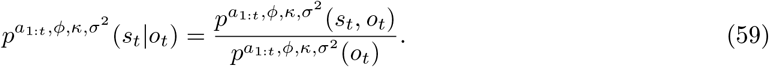

To this end, we first evaluate the marginal distributions

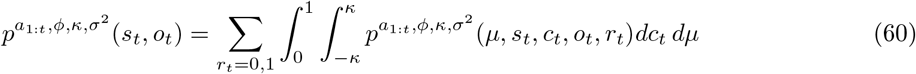

and

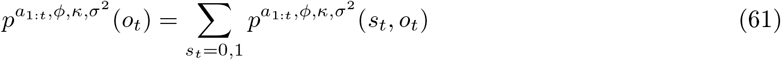

and show that these are not dependent on actions *a*_1:*t*_. We then evaluate (59) and find that the closed-form solution of (7) is also not dependent on *ϕ*.

*Evaluation of* (60)

We first consider the case *a*_*t*_ = 0. We have

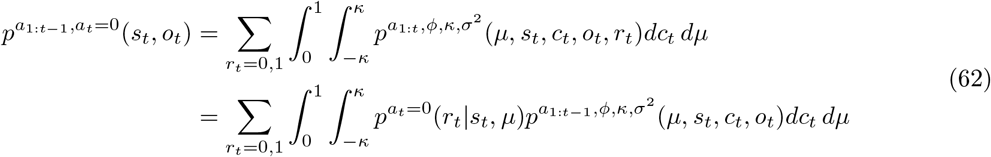

given the conditional independence of *r*_*t*_ of the remaining random variables as defined in (50) and the fact that 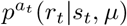 is not parameterized by *a*_1:*t−*1_. Substitution of the functional form (4) of 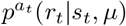 for *a*_*t*_ = 0 then yields

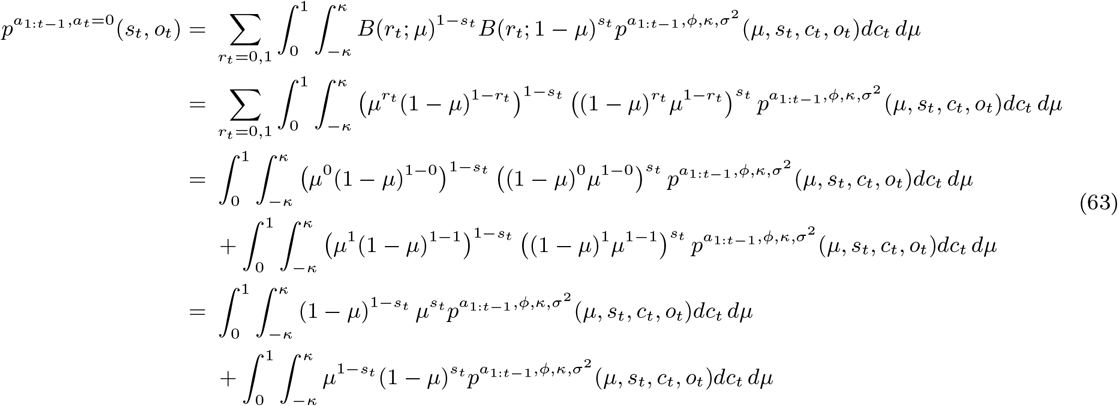

Likewise, substitution of 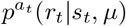 for *a*_*t*_ = 1 yields

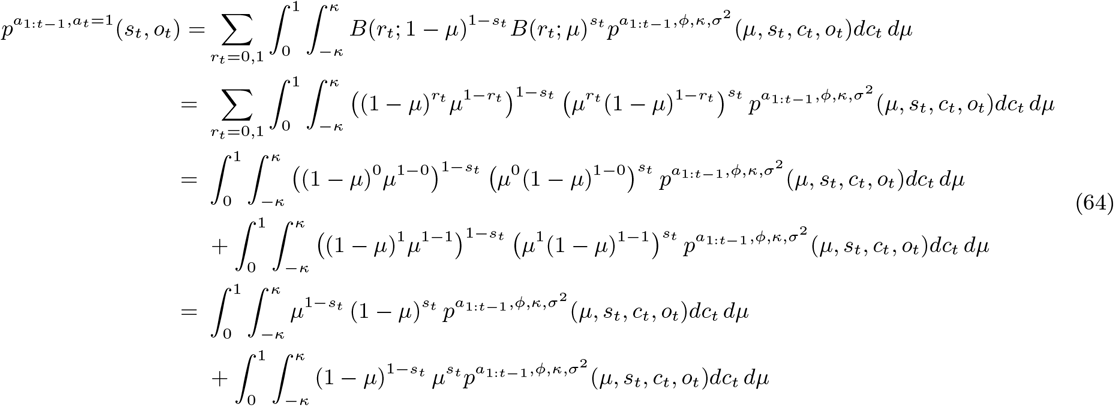

We thus find

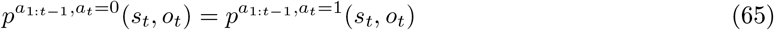

This also entails that the marginal distributions (61), and hence also the belief-state distributions (59) are identical for *a*_*t*_ = 0 and *a*_*t*_ = 1 and thus not dependent on the value of *a*_*t*_. We thus continue the evaluation of the joint distribution of states and observations irrespective of the parameter *a*_*t*_ from (63):

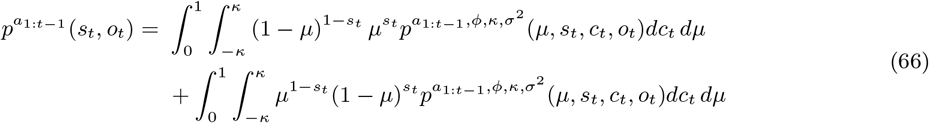

The conditional independence properties and parameter (non)dependencies of the distribution 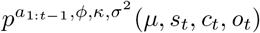 (cf. eq. (50)) then yield

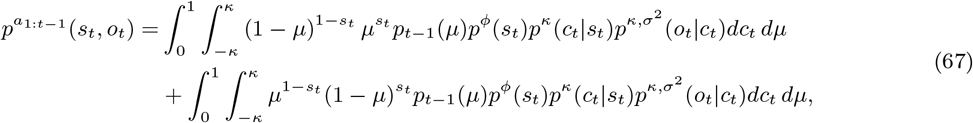

where the dependency on actions *a*_1:*t−*1_ is subsumed in *p_t−_*_1_(*μ*) (cf. eq. (51)). We hence have

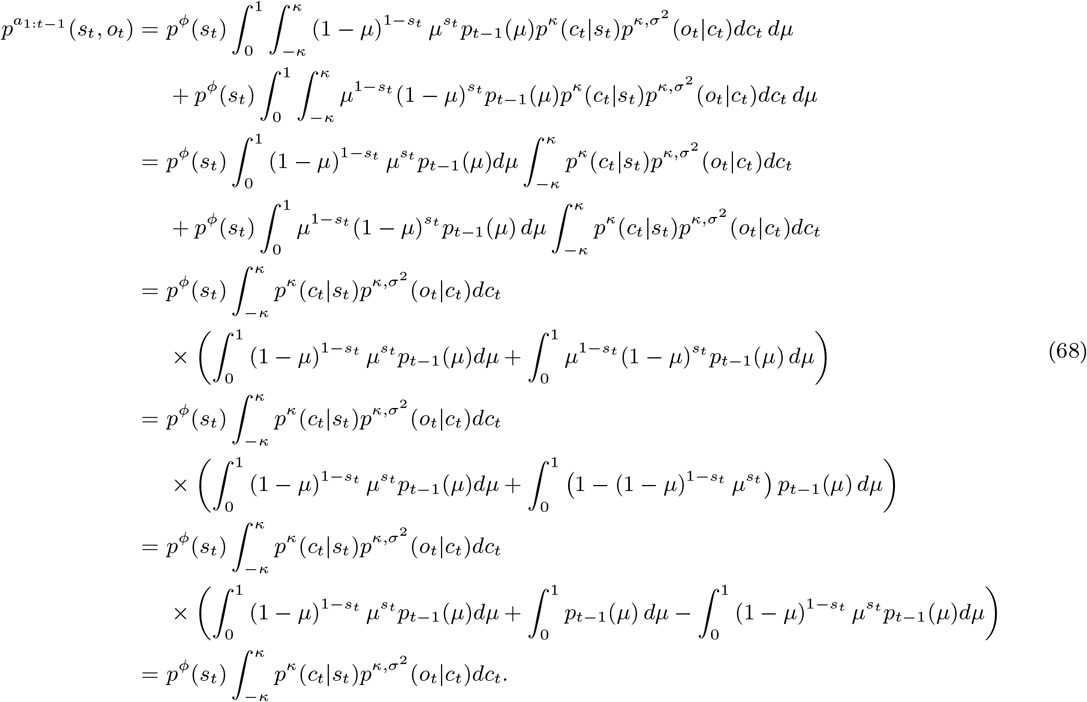

In the fourth equality of the above, we used the fact that

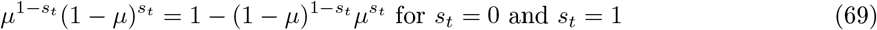

and in the last equality, we used the fact that the contingency parameter marginal distribution *p*_*t*_(*μ*) is represented by a probability density function that integrates to 1 on the support set [0, 1] of *μ*. Note that (68) entails that the belief-state distribution is independent of the contingency parameter marginal distribution, and hence the history of actions *a*_1:*t−*1_. We are thus led to consider the remaining integral explicitly. Substituting the functional forms of the respective probability distributions (cf. (2), (3), (54), (55), (56)), we have

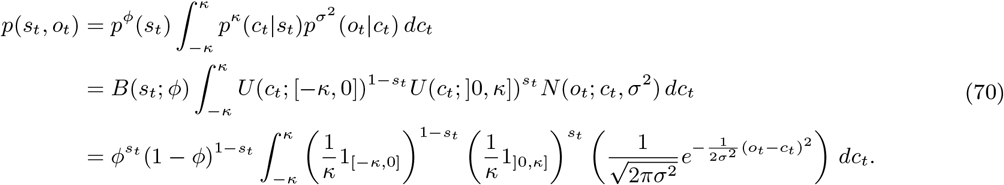

Because either *s*_*t*_ = 0 or *s*_*t*_ = 1, we then have

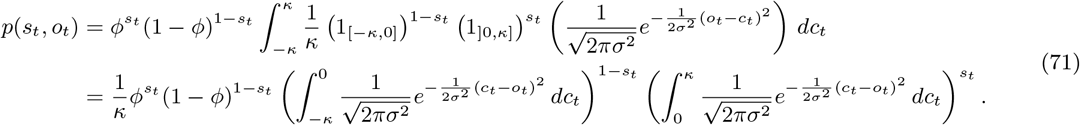

Finally, with the definition of the cumulative density function of the Gaussian distribution (57), we then have

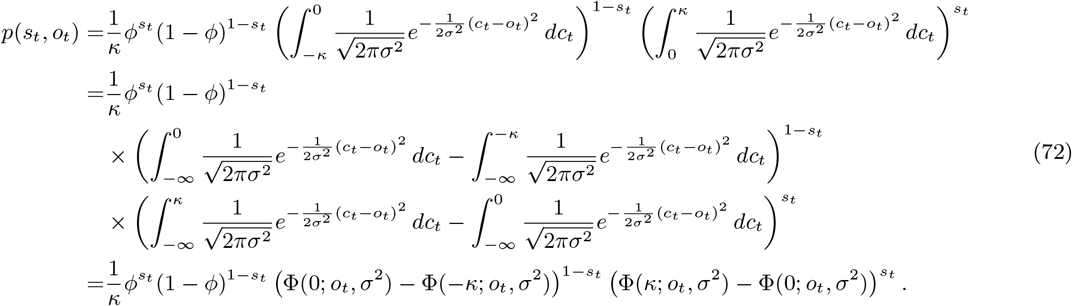

Reintroducing the parameter dependency on the left-hand side of the above, we thus find

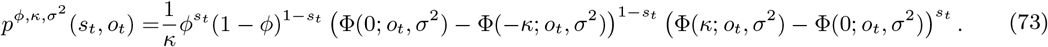

*Evaluation of* (61)

Continuing from (73), we have

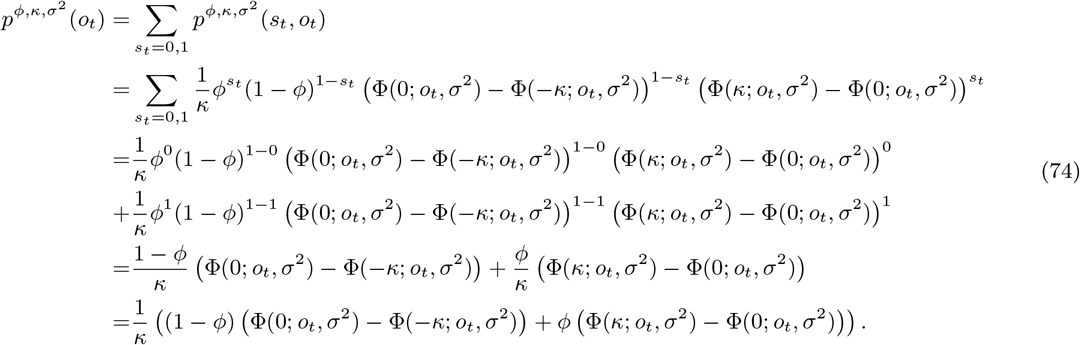

*Evaluation of* (59)

Based on (73) and (74), we have

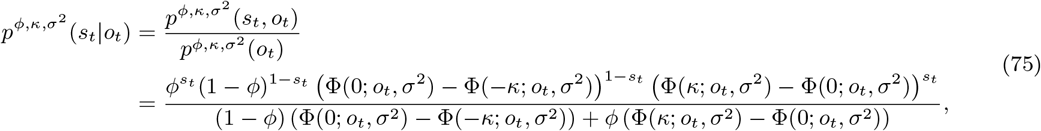

which is the general belief state for arbitrary values of *ϕ ∈* [0, 1]. Using the experimental definition of *ϕ* ≔ 0.5 and thus 1 *− ϕ* = *ϕ*, we obtain

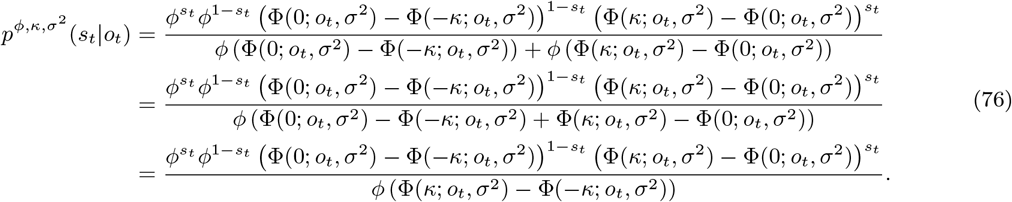

Finally, because either *s*_*t*_ = 0 or *s*_*t*_ = 1, we obtain

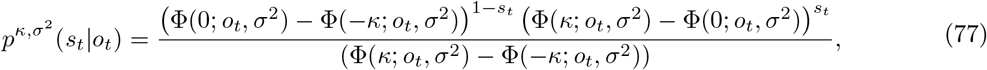

which corresponds to the closed-form solution for the belief state reported in eq. (7).

## 1.2 Proof of(53)

The probabilistic model of interest to compute the observation- and reward-conditional reward probability corresponds to

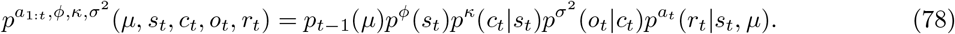

Equation (53) then follows directly from the respective conditional independence properties. For ease of notation, we omit all parameter dependencies on the respective distributions in the following, except for the dependency on *a*_1:*t*_. The observation- and reward-conditional reward distribution can be expressed as

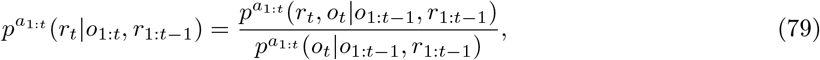

which we compute by first evaluating

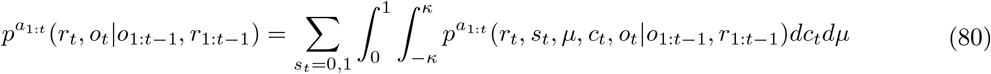

and

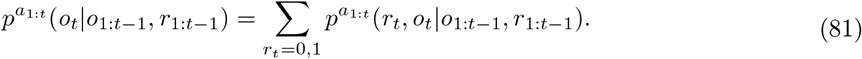

*Evaluation of* (80)

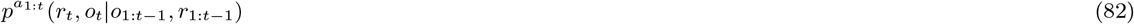

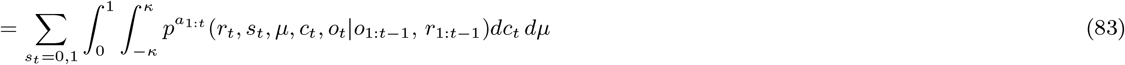

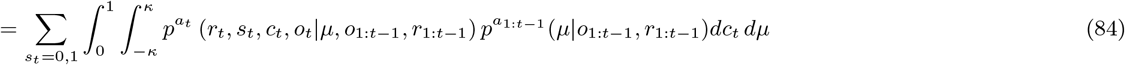

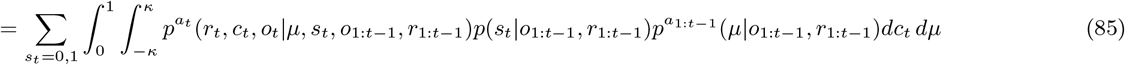

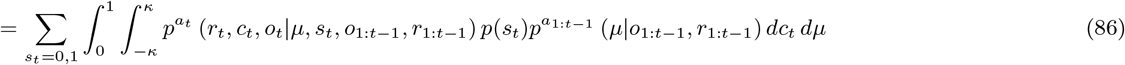

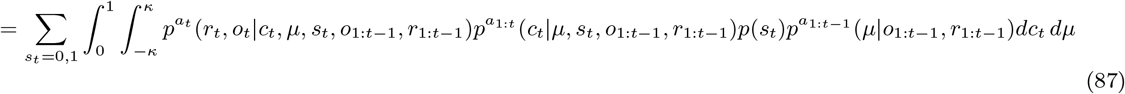

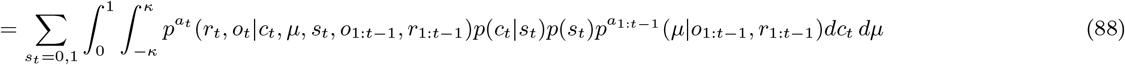

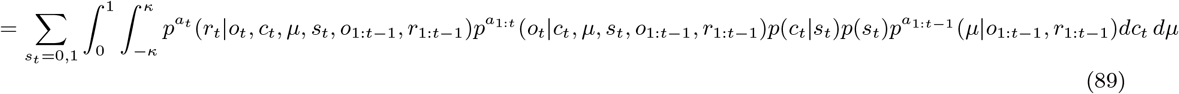

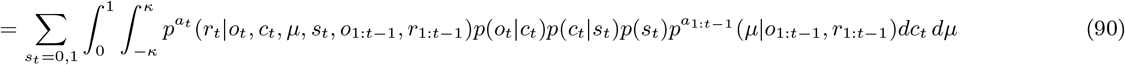

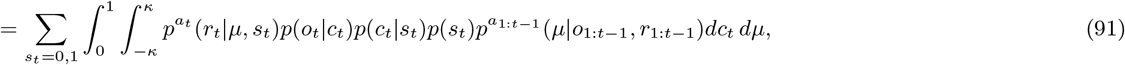

where, in an alternating fashion, we again used the conditional independence properties of the agent model and the calculus of conditional probabilities.

*Evaluation of* (81)

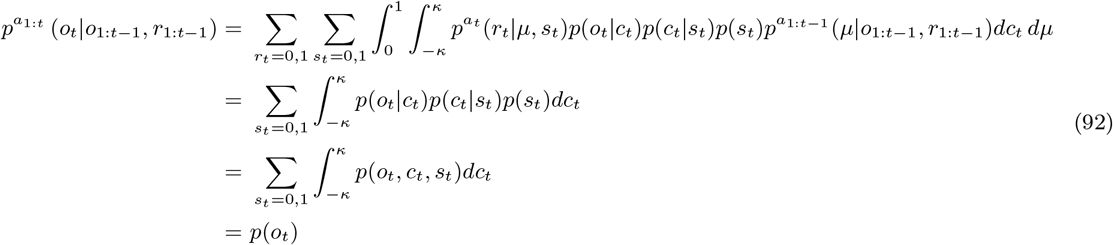

*Evaluation of* (79)

To evaluate eq. (79), we integrate over *c*_*t*_ and render the expression explicitly dependent on the belief state:

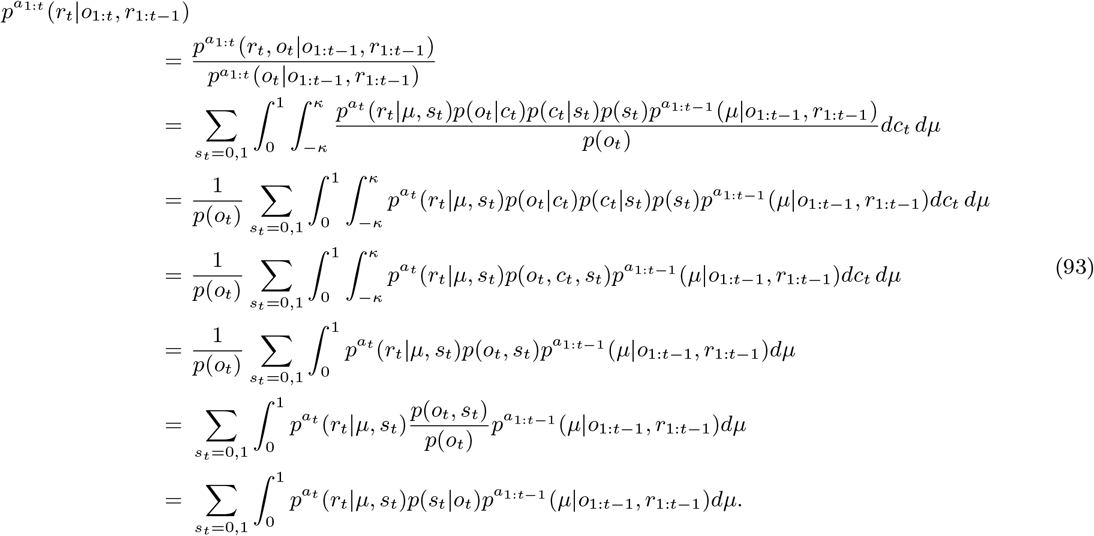

Reintroducing the parameter-dependency in the last equation of (93) for the experimental parameter *ϕ* = 0.5 then yields (53):

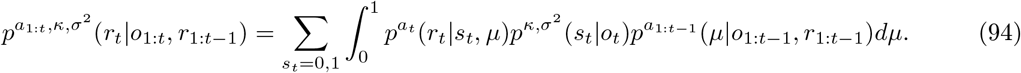

## 1.3 Proof of(11) and (12)

By definition, we have

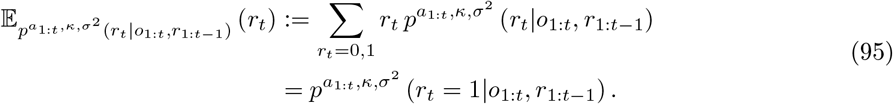

With (53), we thus obtain

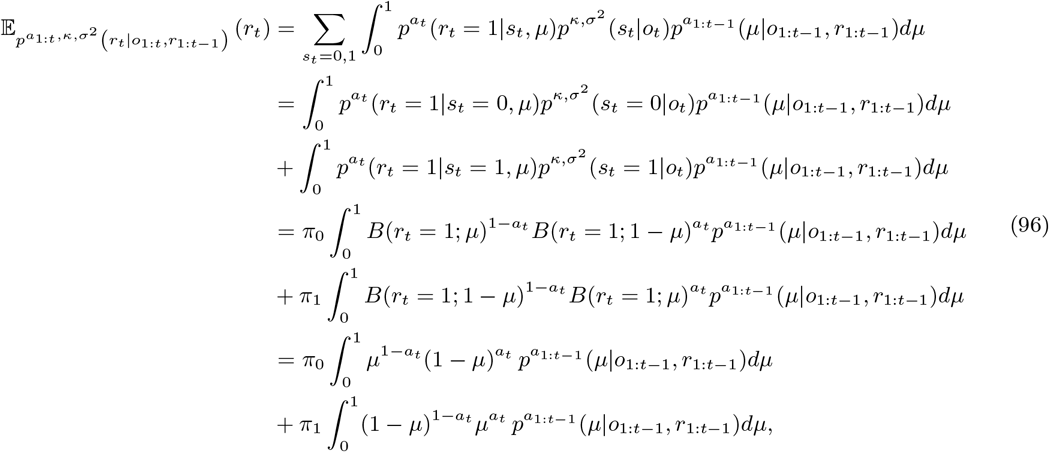

where for the penultimate equality we used the definitions (cf. (7))

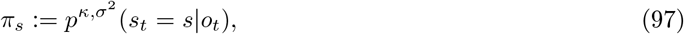

and (cf. (4))

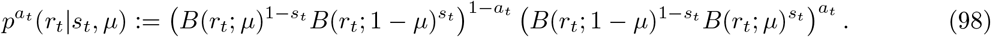

We are finally required to consider the cases *a*_*t*_ = 0 and *a*_*t*_ = 1.

- Case *a*_*t*_ = 0. In this case, we have with (96)

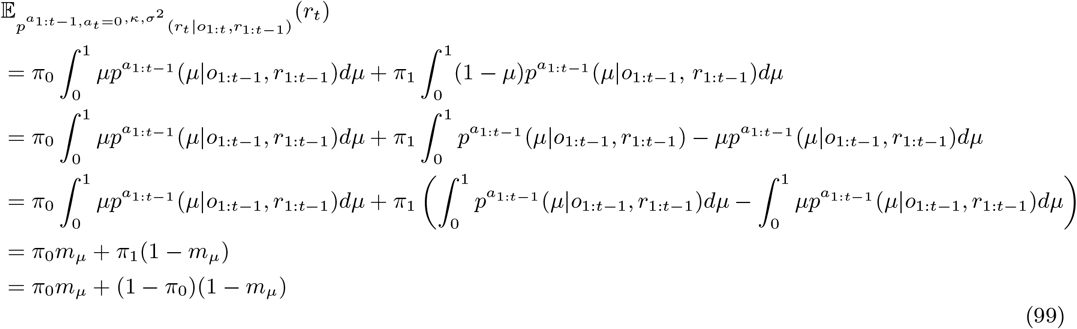

where we used the definition (cf. (13))

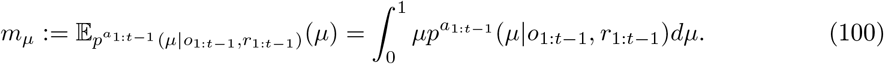

- *a*_*t*_ = 1. In this case, we have, similarly to the above,

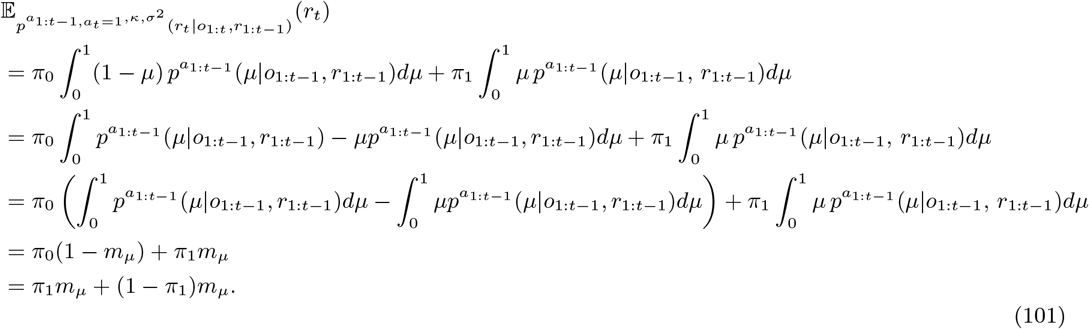

## 1.4 Proof of (14)

The probabilistic model to estimate the contingency parameter corresponds to

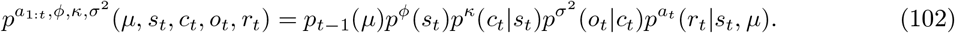

To proof that (14) holds, we derive the parameter distribution

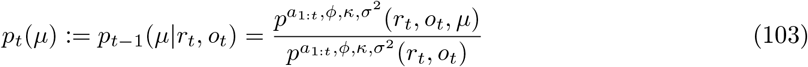

based on an evaluation of the marginal distributions

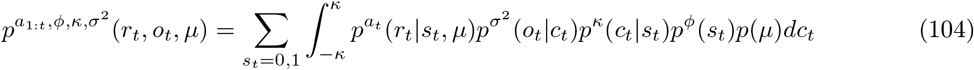

and

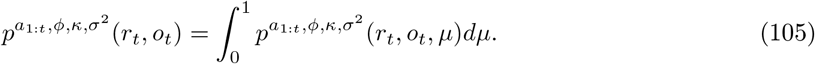

*Evaluation of* (104)

We begin with a substitution of the functional form of 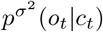 and *p^κ^*(*c*_*t*_ |*s*_*t*_) (cf. eqs (3), (6)).

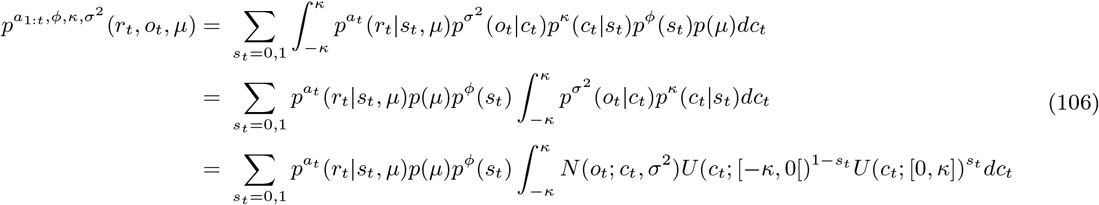

A simplification then yields (cf. eqs. (54), (56))

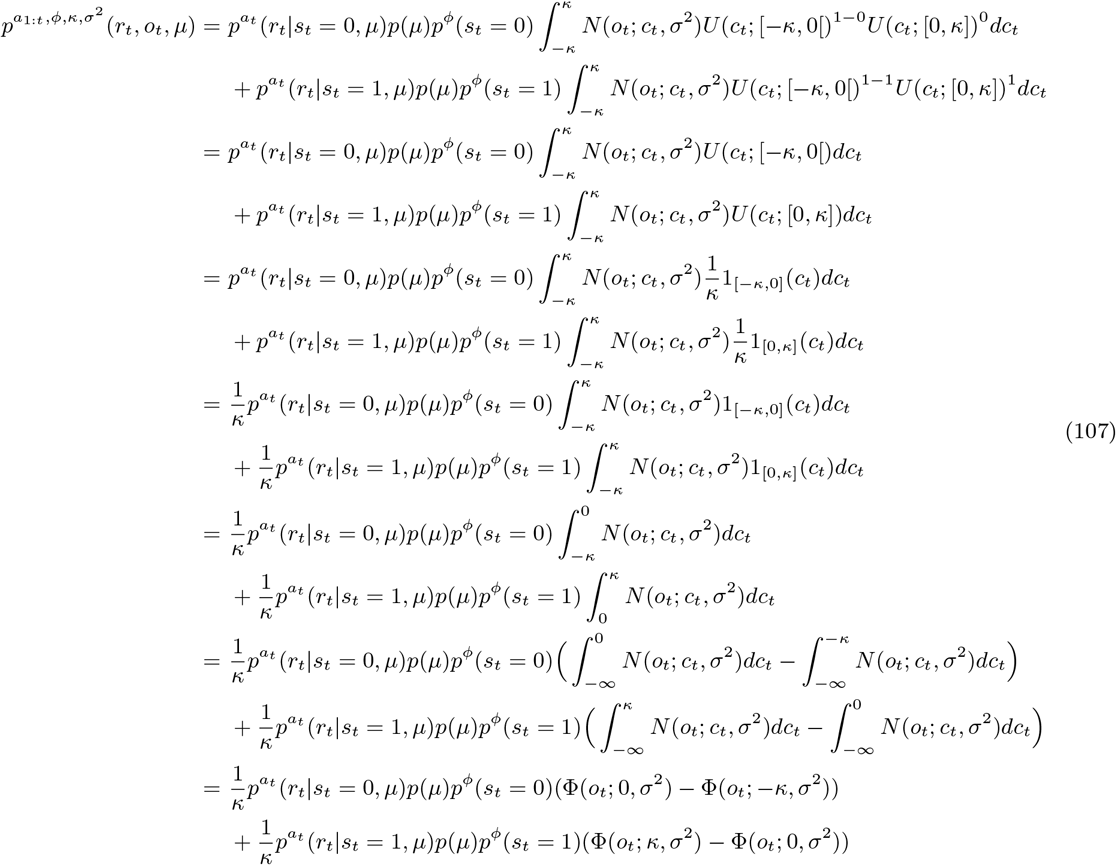

We continue by substituting the functional form of 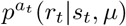 (cf. (4)), *p*(*μ*) (cf. section Bayesian inference agents A1-A3) and *p^ϕ^*(*s*_*t*_) (cf. (2)) and further simplifications

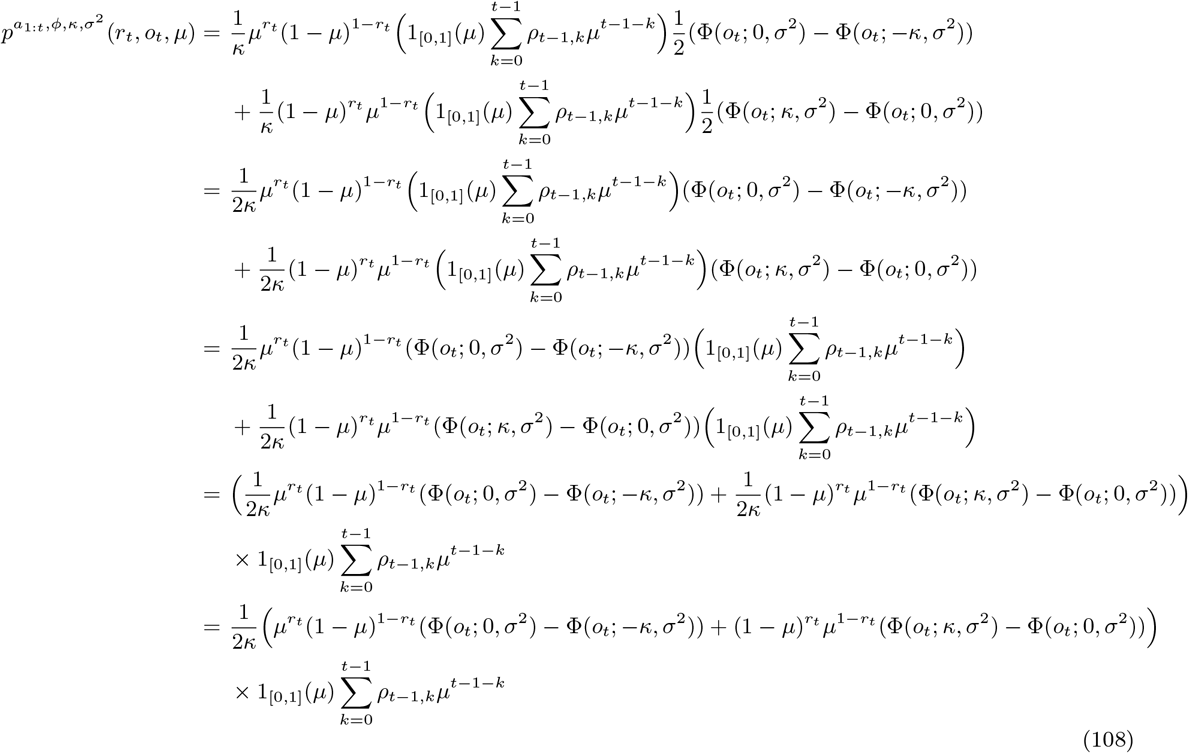

*Evaluation of* (105)

We first rearrange 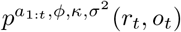 in order to group together terms that are affected by the integration over *μ*.

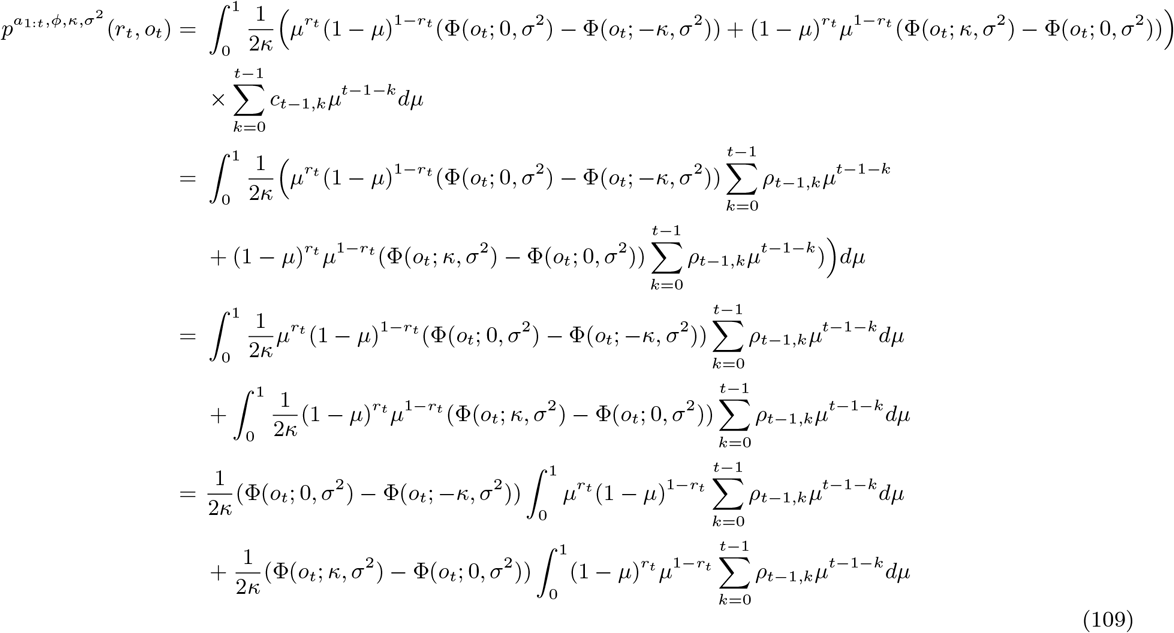

Based on this, we now rearrange these terms to more easily integrate over *μ*.

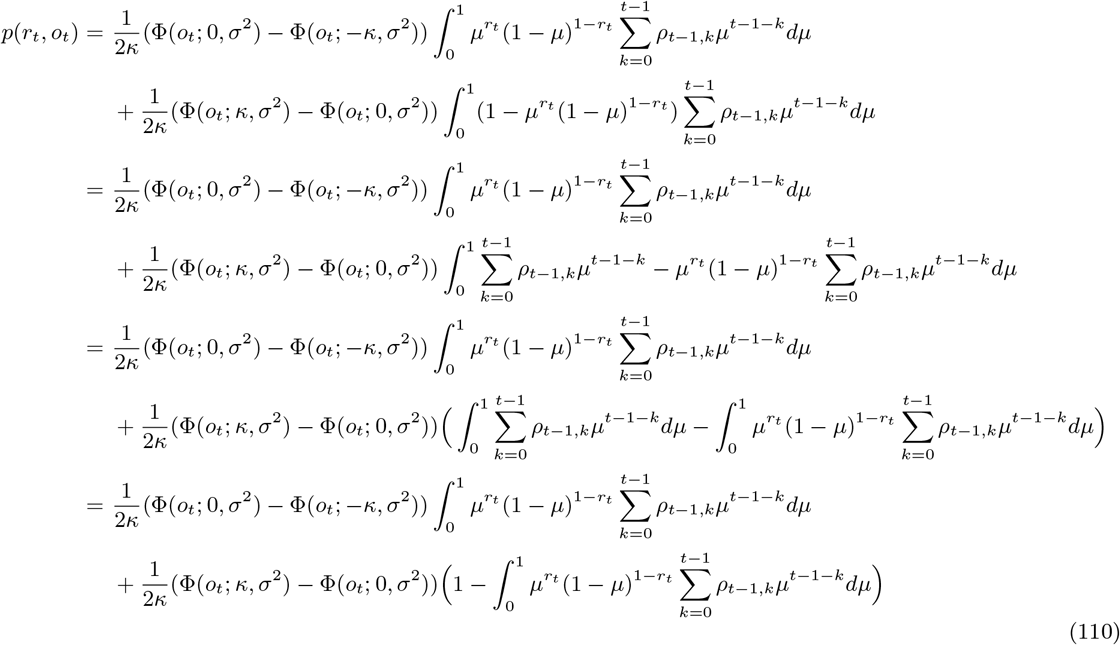

Next, the term 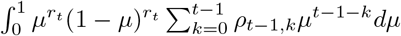 can further be simplified. For *r*_*t*_ = 0, we have

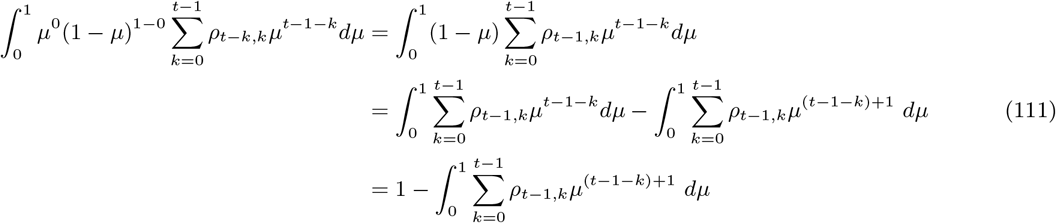

Similarly, for *r*_*t*_ = 1 we have

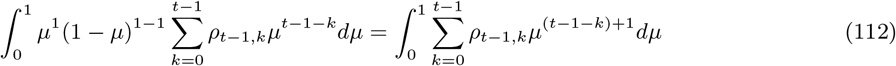

Hence

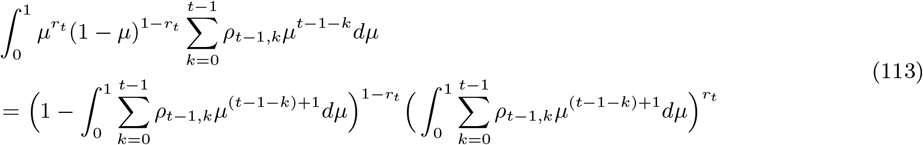

With the definition

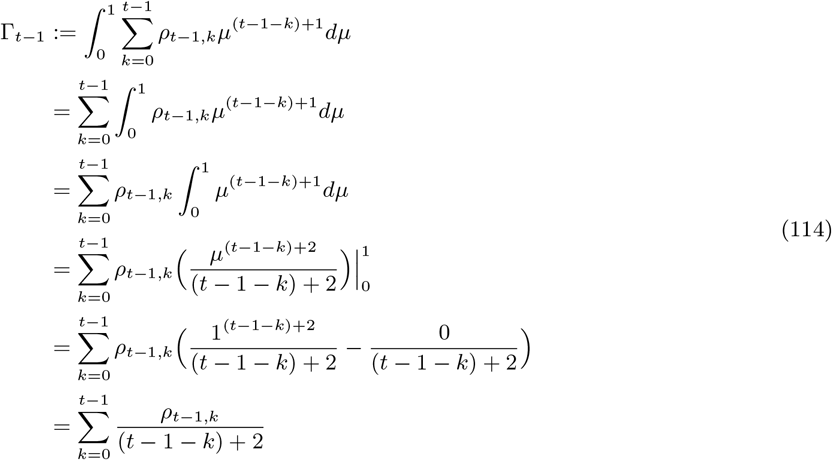

and

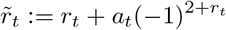

(cf. eq. (18)) we thus have

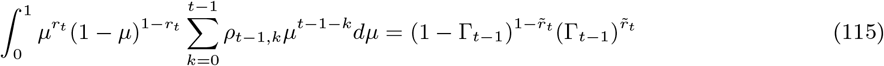

Continuing with eq. (110), we hence obtain

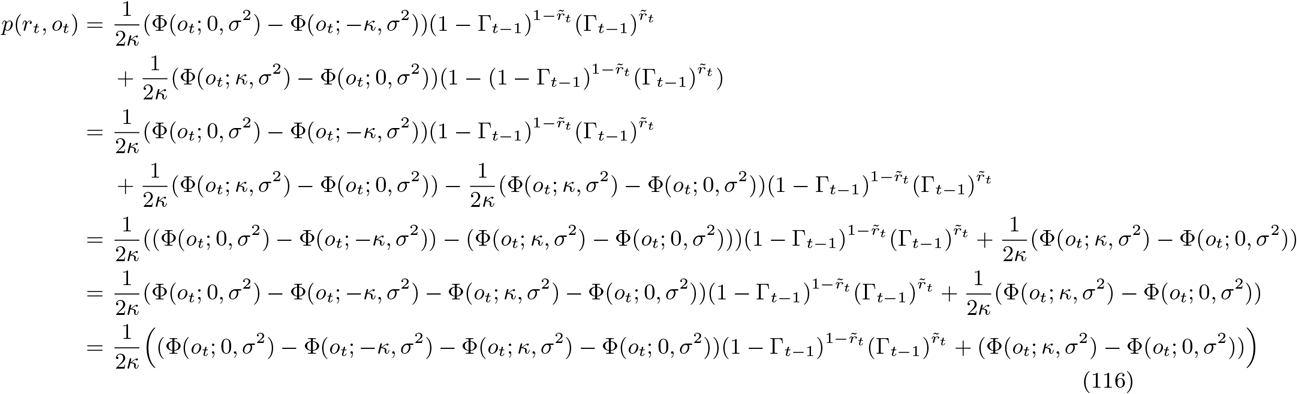

*Evaluation of* (103)

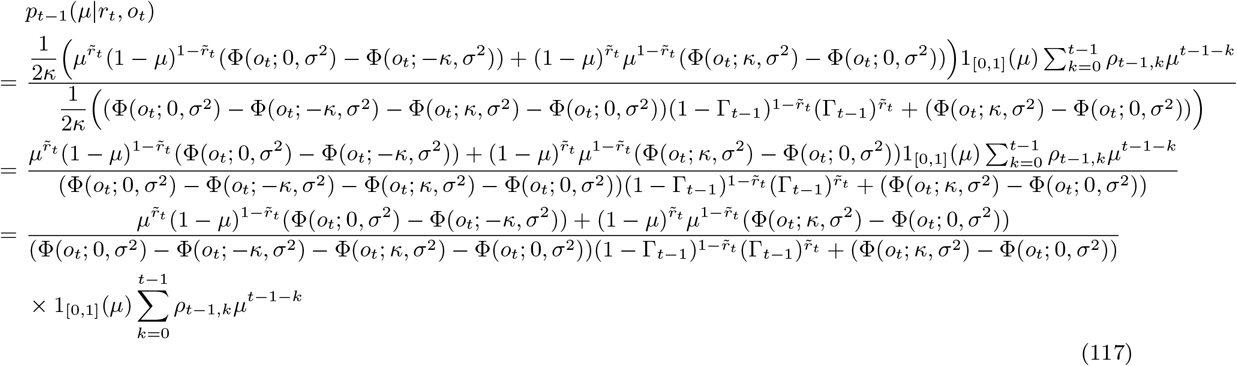

Continuing with the fraction in eq. (117) then yields

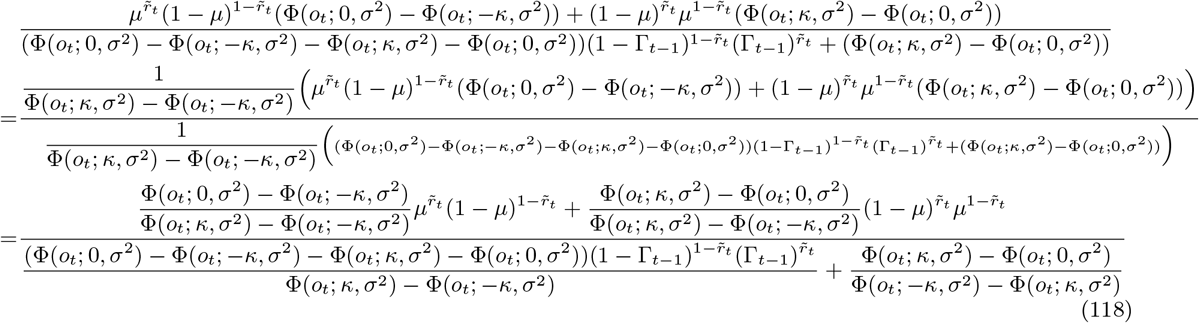

Defining 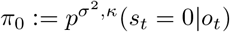 and 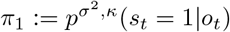 then yields

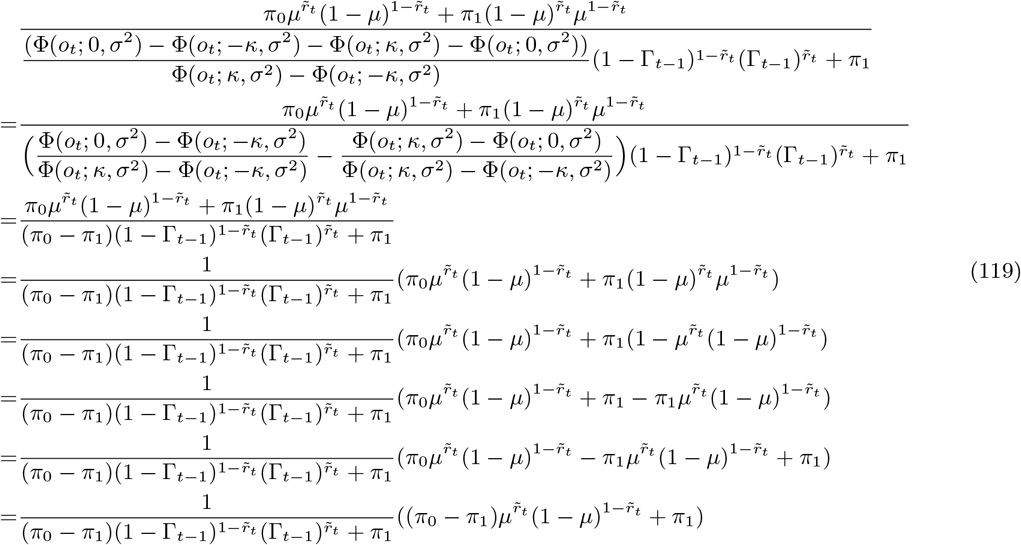

Thus, in combination with eq. (117), we have

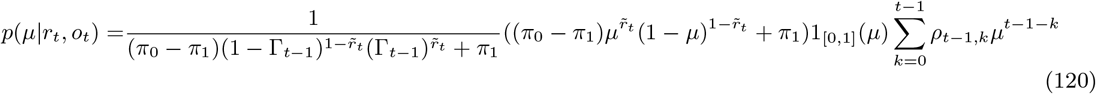

To further simplify the equation, we next show that

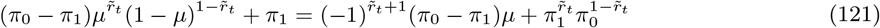

Case 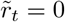

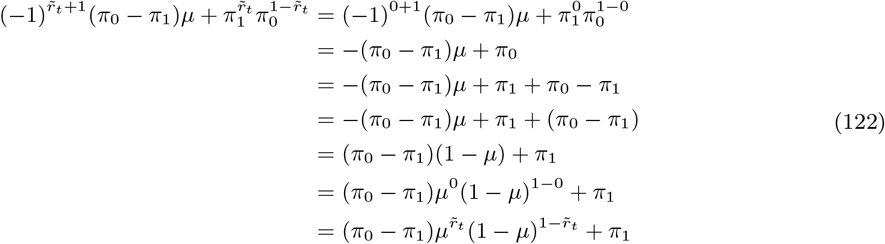

Case 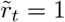

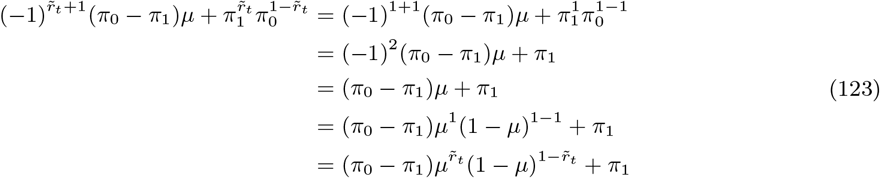

This yields

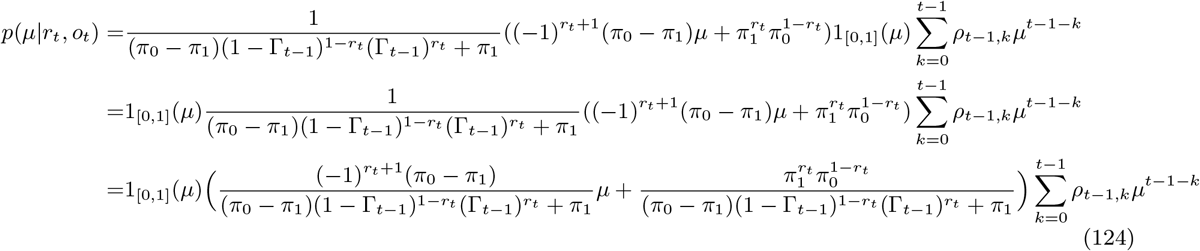

We next define

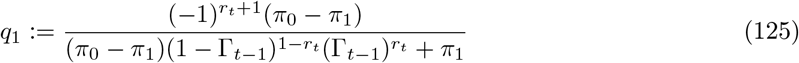

and

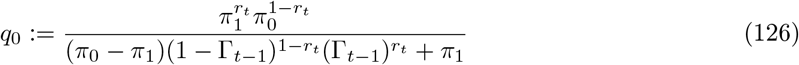

and thus obtain

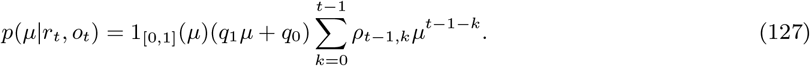

## 1.5 Proof of (31)

For this case, the marginal probabilistic model of interest corresponds to

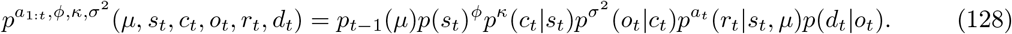

To proof that eq. (31) holds, we derive the contrast-difference-conditional perceptual-choice distribution

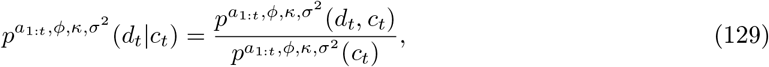

which requires an evaluation of the marginal distributions

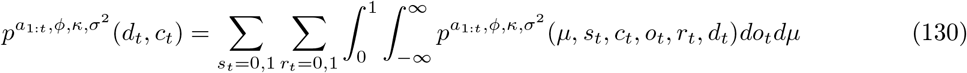

and

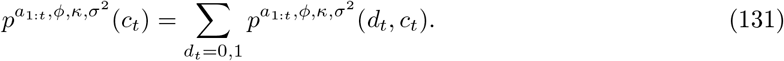

*Evaluation of* (130)

For the case *a*_*t*_ = 0, we have

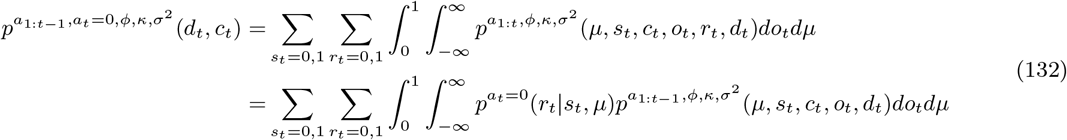

which, as also argued in 1.1, follows from the conditional independence of *r_t_* on the remaining variables and actions *a*_1:*t−*1_. This yields

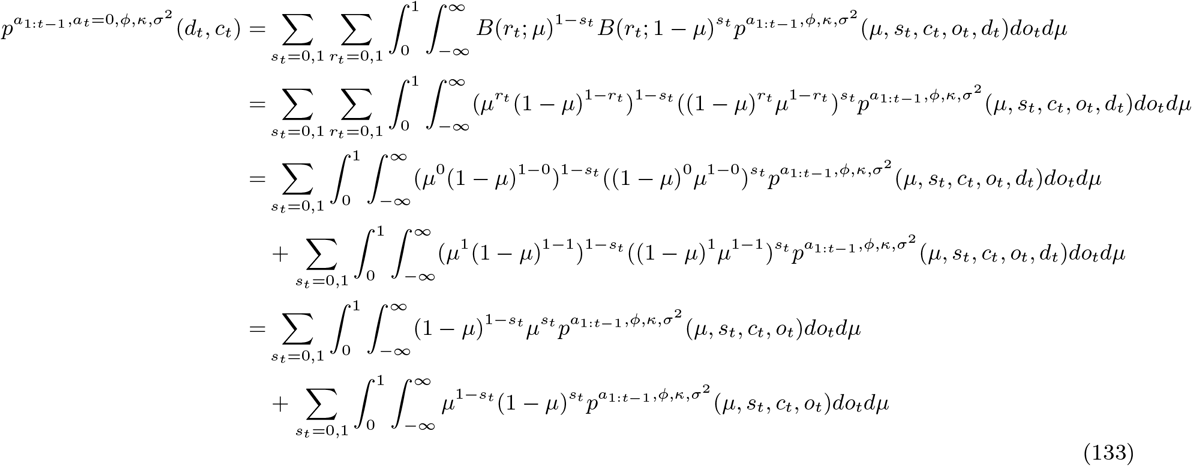

Likewise, for *a*_*t*_ = 1 we obtain

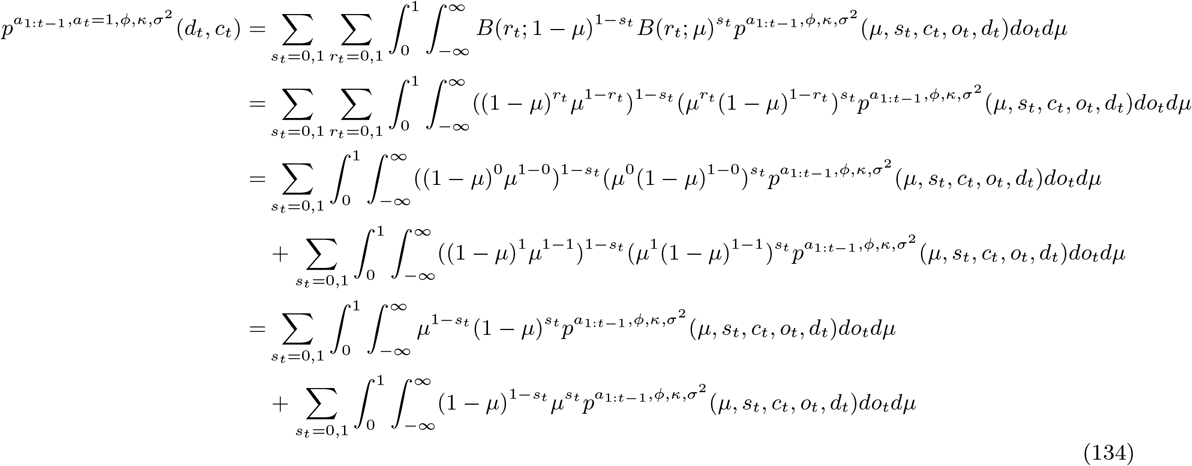

Similar to 1.1, this thus reveals that 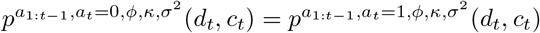 and it follows that the marginal distribution of *d*_*t*_ and *c*_*t*_ is not dependent on action *a*_*t*_. We thus continue with

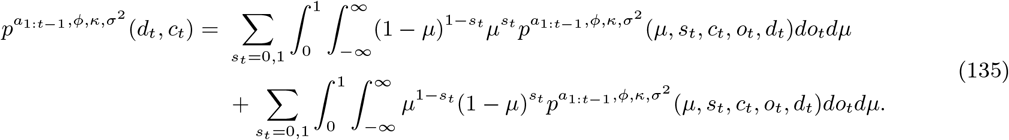

Based on the conditional independence properties and parameter dependencies shown in eq. 128, we can rewrite eq. 135 as (cf. eq. 68)

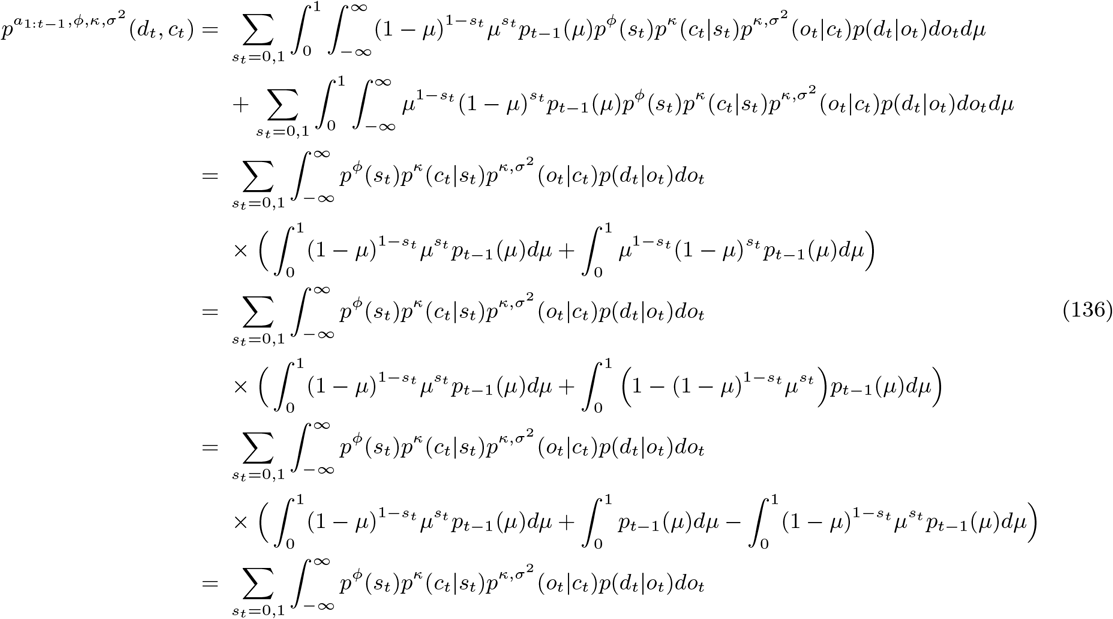

We next define the observation-conditional perceptual-choice probability

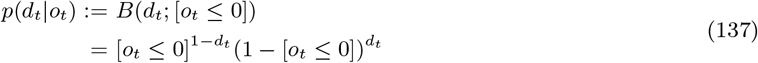

where we use the Bernoulli distribution with a Bernoulli parameter encoded by the the Iverson bracket defined as

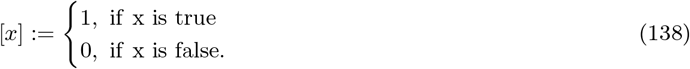

This thus reflects that the agent deterministically chooses *d*_*t*_ = 0 if *o*_*t*_ ≤ 0 and *d*_*t*_ = 1 if *o_t_ >* 0, which follows from our definition of the agent’s perceptual-choice policy (eq. 8) and *ϕ* ≔ 0.5, indicating equal prior state probabilities. In functional form, eq. (136) thus corresponds to

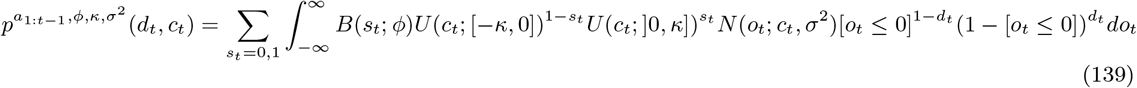

Evaluating (139) then yields

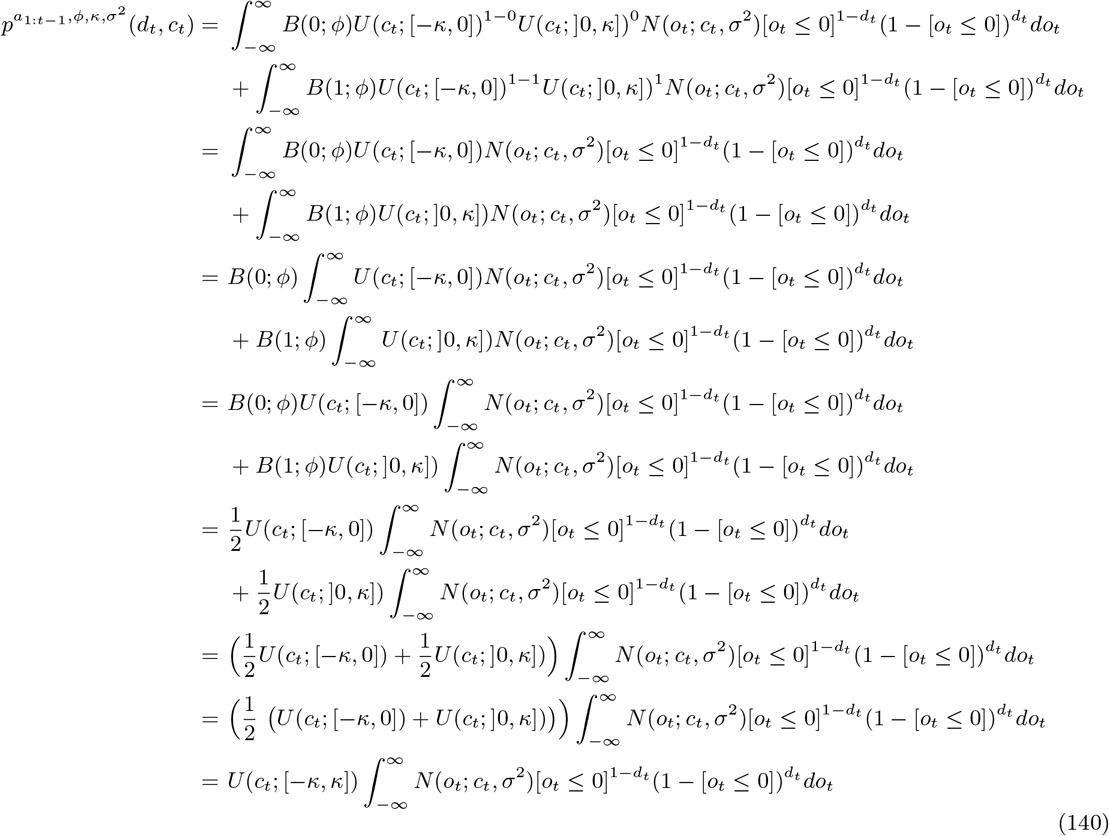

Based on the conditional dependence of the agent’s perceptual decision on [*o*_*t*_ ≤ 0] as defined above, we can split the integral as follows

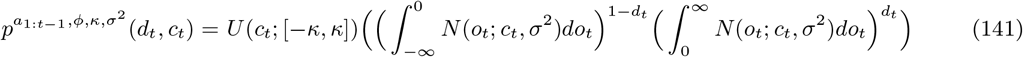

Finally, we can express the integrals over observations as Gaussian CDF’s

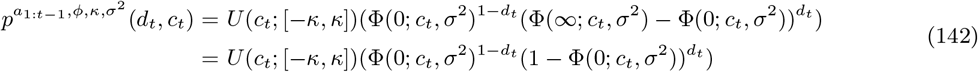

**Figure SM 6.**
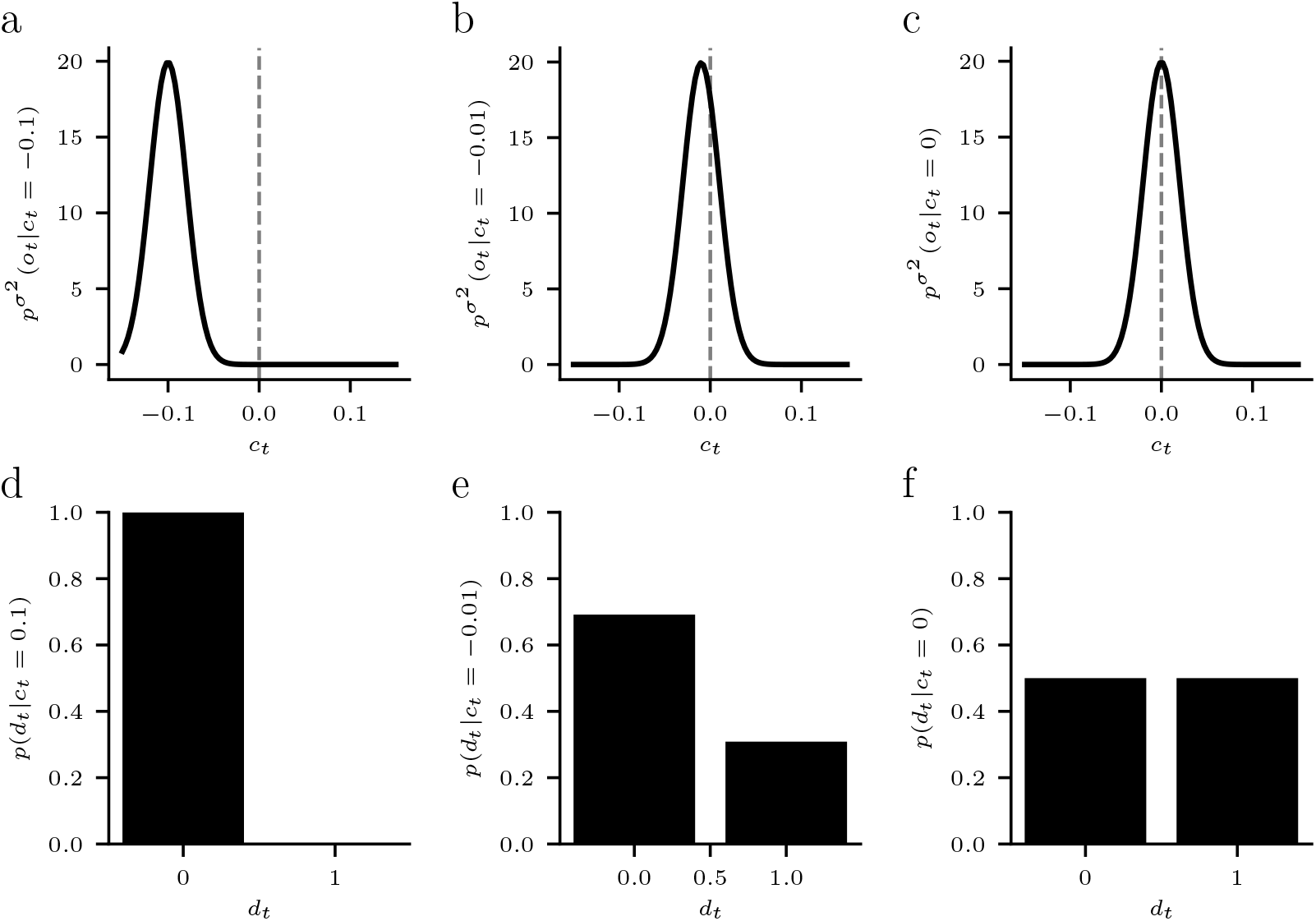
Probability of a perceptual decision for different contrast differences. **a-c)** Gaussian probability density function over observations conditional on different contrast differences. **d-f)** Corresponding probability for perceptual decision conditional on contrast differences.

*Evaluation of* (131)

Continuing from eq. (142), we have

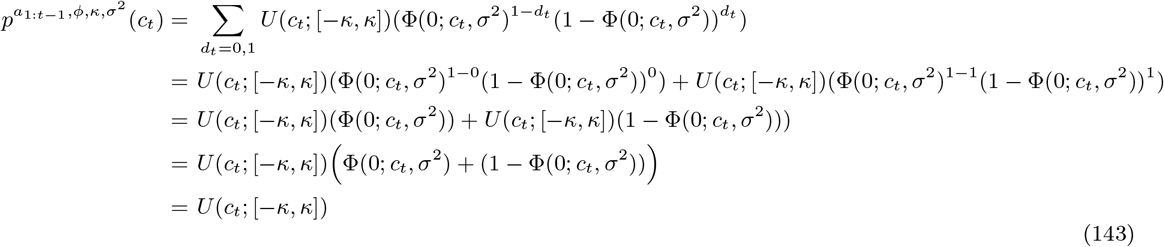

*Evaluation of* (129)

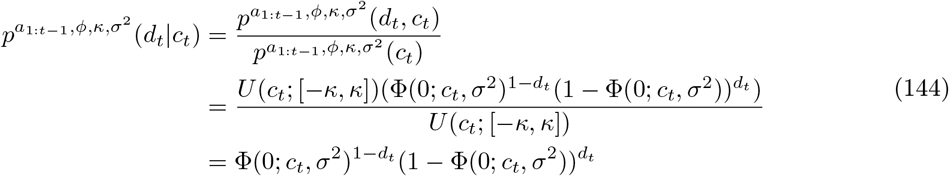

In Figure SM 6, we plot an example of the relation between *p*(*o_t_|c_t_*) and *p*(*d_t_|c_t_*) for different levels of *c*_*t*_.

## Footnotes

1 https://www.psychopy.org/api/visual/gratingstim.html

